# Friend and Foe: Genome-Wide Analysis of the Tardigrade Dsup Protein Expressed in Yeast Reveals Trade-offs of DNA Protection

**DOI:** 10.64898/2025.12.24.696340

**Authors:** Hamid Kian Gaikani, Marjan Barazandeh, Lewis Hitchens, Guri Giaever, Corey Nislow

**Affiliations:** Faculty of Pharmaceutical Sciences, University of British Columbia, Vancouver, BC, Canada; Department of Chemistry, University of British Columbia, Vancouver, BC, Canada

**Keywords:** Dsup, yeast, tardigrade, DNA damage, Yeast Knockout Collection, genome maintenance, synthetic lethality, oxidative stress, MMS, bleomycin

## Abstract

Tardigrades are among the most resilient eukaryotes known, capable of surviving a wide range of extreme conditions. While their genomes contain thousands of tardigrade-specific genes, one key contributor to their stress resistance is the Damage suppressor (Dsup) protein. Previous work showed that expression of Dsup in cancer cells reduced DNA damage caused by X-rays. This observation has sparked growing interest in Dsup across diverse model organisms. Most studies to date have focused on phenotypic outcomes, while the mechanisms and consequences of Dsup remain poorly understood. Deciphering its mechanism is crucial, especially because Dsup expression is deleterious to some cells and organisms. In this work, we show that Dsup protects yeast against a variety of DNA-damaging agents. However, this protection is not universal, and it has a fitness cost. Analysis of nucleosome occupancy of cells expressing Dsup shows that effects on chromatin are modest, without global remodeling, and transcriptome analysis reveals a robust induction of oxidative stress pathways, which might be a form of preconditioning for oxidative stress. Genome-wide screens of ∼4300 loss-of-function mutants expressing Dsup shows that deletions incompatible with Dsup are highly enriched for DNA repair processes and that under DNA damage stress, Dsup can protect some mutants in a pathway-specific manner. Our findings demonstrate that while Dsup broadly enhances resistance to genotoxic stress, it also imposes physiological costs and generates new dependencies on DNA repair and genome maintenance pathways.

## 1. Introduction

Tardigrades are eukaryotes known for their exceptional resilience to extreme stressors, including desiccation, temperature extremes, oxidative stress, and intense radiation (*1–4*). For instance, *Ramazzottius varieornatus* can survive more than 48 h of acute exposures of 4,000 Gy of gamma radiation (*5*), orders of magnitude higher than the median lethal dose for whole-body human exposure (LD_50_ of ∼4.1 Gy) (*6*). An underlying factor in their radiotolerance is the Dsup, a chromatin-associated, tardigrade-specific disordered protein first discovered by Hashimoto *et al.* (*5*). When Dsup was expressed in human cancer cells, it suppressed X-ray-induced DNA breaks by ∼40% and improved cell survival after irradiation (*5*).

In vitro assays show Dsup can coat nucleosomes and reduce chromatin cleavage by hydroxyl radicals (*7*). Whether this protection arises purely from physical shielding or if it also involves specific cellular functions remains unresolved. Notably, some studies have reported Dsup-dependent upregulation of DNA repair genes (*8*), suggesting a potential role beyond passive protection. Although Dsup has attracted widespread interest as a tool to enhance stress resistance in other organisms, its transgenic expression is not completely benign. While the initial study in human HEK293 cells showed that Dsup reduced DNA damage without an obvious effect on cell fitness (*5*), recent investigations in other organisms and cell types reveal a more complex, context-dependent picture. A study in *C. elegans* showed that expressing Dsup enhances resistance to oxidative and radiation stress, largely by reducing ROS levels and suppressing mitochondrial respiration without affecting development or fertility (*9*). On the other hand, in transgenic flies, expression of Dsup led to enhanced survival after gamma irradiation and oxidative stress, yet it also caused widespread transcriptional repression and disrupted locomotor activity (*10*). In mammalian neurons, Dsup expression caused chromatin condensation, induced DNA double-strand breaks, and reduced cell viability (*11*). These findings demonstrate that transgenic Dsup affects different hosts differently and emphasize the importance of cell type when interpreting Dsup’s function. In another recent study, yeast cells expressing Dsup showed enhanced survival under chronic oxidative DNA damage, yet Dsup afforded no protection, and even increased sensitivity to MMS, bleomycin, and UV (*12*). One hypothesis to explain these discordant observations is that in foreign cells, Dsup could impose novel stresses or dependencies on existing pathways. Yet the specific cellular pathways involved are unknown.

In this study, we sought to address these knowledge gaps by leveraging yeast as a eukaryotic model that offers powerful, unbiased screening tools to explore genetic interactions on a genome-wide scale. We expressed Dsup in yeast and examined its influence on cellular physiology and stress response, while systematically mapping genetic dependencies for Dsup via synthetic lethal screens. Contrary to the aforementioned study in yeast (*12*), we found that Dsup expression in yeast improves survival under a variety of genotoxic stresses. Nevertheless, we also observed quantifiable physiological costs. For example, Dsup-expressing yeast grows more slowly and exhibits a cell-cycle delay. Genome-wide screening revealed that Dsup creates new genetic liabilities towards DNA repair and genome maintenance pathways, as well as in cell cycle regulators. We also found that Dsup can rescue certain deletions that are hyper-sensitive under certain conditions, suggesting its protection is pathway-specific. Together, our results paint a comprehensive picture of Dsup’s double-edged nature in a eukaryotic cell. On one hand, Dsup shields DNA, reducing damage and enhancing survival. On the other hand, Dsup perturbs replication and transcription and amplifies the cell’s dependencies on genome-maintenance processes. This genome-wide study of Dsup’s effects not only demonstrates the protein’s protective benefits and its deleterious side-effects but also shows critical pathways and trade-offs to consider when repurposing extremophile proteins for synthetic biology or biotechnology. By examining how Dsup interacts with a model eukaryotic genome, this study provides insight into the feasibility and limitations of engineering stress tolerance across species.

## 2. Results

### 2.1 Heterologous expression of Dsup in yeast is not benign and cells silence centromeric plasmids

To express the Dsup protein in haploid yeast cells (BY4741), we cloned a codon optimized gene with a GFP tag at the C-terminus into a centromeric plasmid under the *TDH3* promoter. 2 days after transformation, single colonies were observed on the plate, but the expression of Dsup-GFP was silenced. In contrast, all the colonies transformed with a control GFP-only plasmid expressed GFP at high levels (Fig. 1 a and b). Next, using a centromeric plasmid containing Dsup-GFP downstream of a *GAL1* promoter, we obtained colonies, but observed that in galactose (inducing) medium, cells carrying the Dsup plasmid stayed in lag phase for ∼40 h before they started doubling. At the end of logarithmic growth, fewer than 3% of Dsup-GFP cells showed detectable fluorescence, compared with a bright, uniform signal in GFP-only controls. Microscopic observation of cultures in galactose medium after 24 hours showed that ∼97% of cells expressing Dsup-GFP were arrested in G2/M, characterized by large, budded morphology (Fig. 1c-f). We therefore tried a CEN6 plasmid with a β-estradiol-inducible promoter, which successfully expressed Dsup. However, fluorescence microscopy showed that the expression was highly heterogeneous in contrast to the uniform expression seen for the GFP control (Fig. S1). Integration of the same construct (pEstradiol-Dsup-GFP) into the genome at *LEU2* locus showed a dose-dependent and homogenous expression of Dsup, which was also durable even after 20 generations (Fig. 1g). This construct was used for all subsequent experiments.

**Figure 1.**
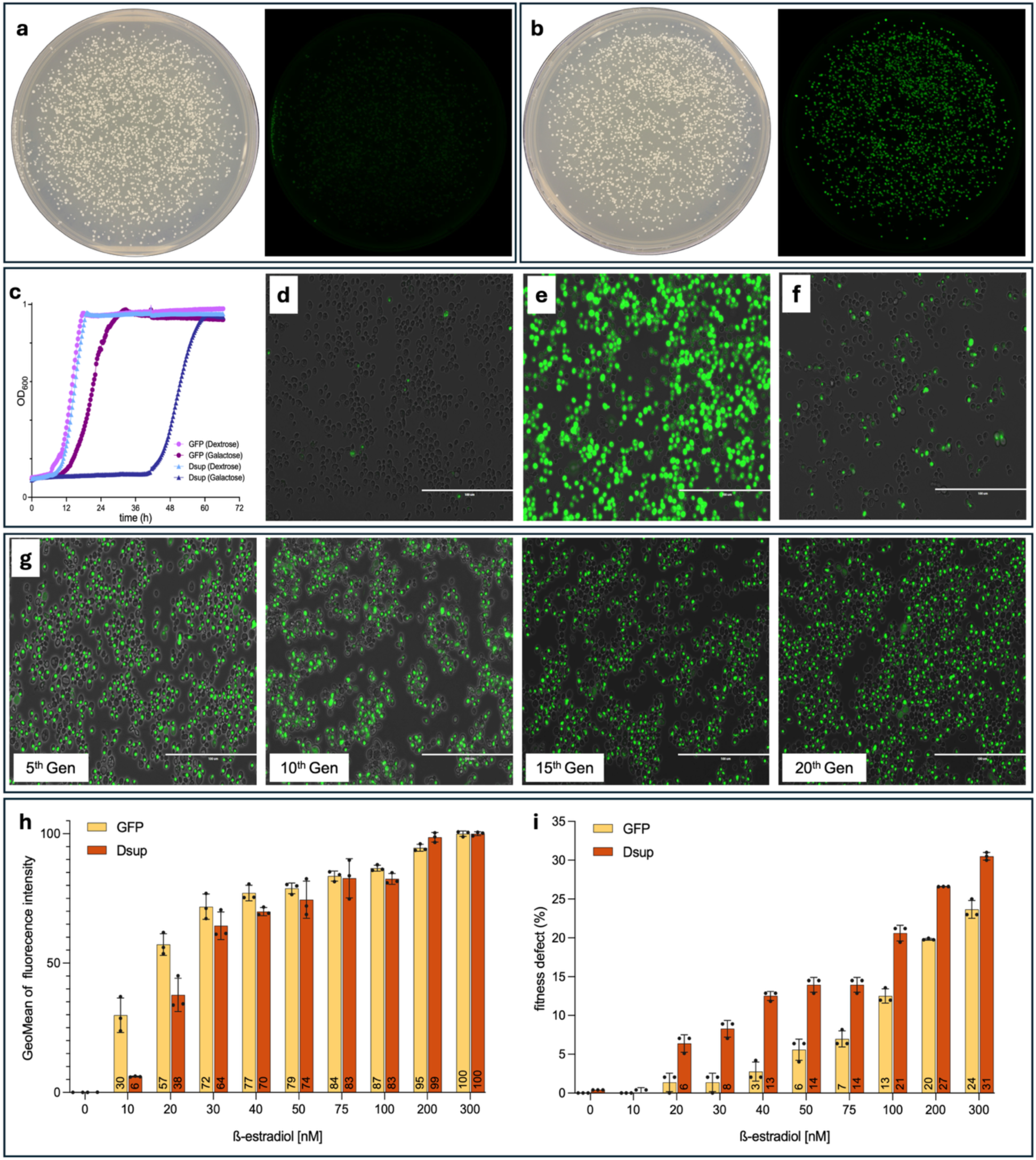
Overview of Dsup expression in yeast. Picture and fluorescence scan of plates of yeast transformed with centromeric plasmids with p*TDH3* Dsup-GFP (**a**) or GFP (**b**) show that while Dsup-GFP expression is silenced and undetectable, all GFP-only colonies have a strong signal. (**c**) Growth curves of strains carrying either GFP or Dsup-GFP plasmids cultured in 2% dextrose or 2% galactose, monitored by optical density at 600nm. (**d**) Dsup-GFP cells at the end of logarithmic phase, and (**e**) GFP-only cells at the end of logarithmic phase (OD ≈ 0.9). (**f**) In cells with p*GAL1*-Dsup-GFP after 24 h in 2% galactose medium, 94 out of 97 (97%) of cells expressing Dsup-GFP are arrested in G2/M. (**g**) Fluorescence images of cells expressing Dsup-GFP shows that expression remains stable after continuous culture for at least 20 generations (scale bar: 100 μm). (**h**) Dose-dependent induction of Dsup-GFP and GFP controls measured by geometric mean fluorescence intensity across a range of β-estradiol concentrations. Values indicate normalized expression levels relative to maximal induction. (**i**) Growth defect caused by Dsup and GFP strains at increasing β-estradiol concentrations. Data represent mean ± s.d. from biological replicates.

Upon induction with increasing amounts of estradiol (from 0–300 nM), Dsup-GFP fluorescence was initially detectable at 10 nM, and the expression was strong and homogeneous at estradiol concentrations of 20 nM and above. This dose-dependent induction plateaued at 75 nM b-estradiol (Fig. 1h), which was selected as the standard induction condition for subsequent experiments. Liquid growth assays showed that expression of both Dsup-GFP and GFP causes a fitness defect, although the fitness defect was worse in Dsup-GFP cells at all concentrations (Fig. 1i).

Fluorescence microscopy showed that Dsup localizes into the nucleus. Given that Dsup binds DNA, we examined its potential localization to mitochondria. Using Cox4–mCherry as a mitochondrial marker, we found no evidence of mitochondrial localization (Fig. S2). Flow cytometry was used to assess the effect of Dsup expression on yeast cell-cycle dynamics. At 0 nM estradiol, most cells were in G1 (64.4%), while 35% were in G2, and approximately 1% were in S phase. A small fraction (0.8%) of cells was detected in S phase. When Dsup was induced with 75 nM estradiol, the G1 population dropped to 52.4%, while the population of cells in G2 increased to 47.4%, and the population of cells in S phase remained very low (0.6%). With stronger induction at 200 nM, the G2 shift became more evident, with the G1 fraction dropping to 37.6%, G2 rising to 52.6%, and the S-phase population increasing to 9.7%. (Fig. S3).

### 2.2 Dsup selectively strengthens transcriptional silencing by increasing nucleosome occupancy

Given its known interaction with nucleosomes, we asked if expression of Dsup influences transcriptional silencing in yeast. We used a yeast strain with three reporter genes integrated at three loci that are subject to transcriptional silencing (the rDNA locus, the silent mating locus HMR, and a telomere) to test whether Dsup affects transcriptional repression *in vivo* (*13*) (Fig. 2a). This experiment gives a semi-quantitative readout—transcriptional silencing and de-repression occur along a spectrum. Hence, the outcome can range from no growth to full growth. In SC medium, both Dsup^−^ and Dsup^+^ cells grew after six dilutions (5x dilution factor). Dsup^+^ colonies were smaller, which reflects the fitness defect caused by Dsup expression. The –Ura and 5-FOA plates measuring Ura3 expression and silencing, respectively, show that most cells express Ura3, able to grow on –Ura medium but fail to grow on 5-FOA. This result shows that Dsup does not alter telomeric expression of the *URA3* marker (Fig. 2b). For the *ADE2* reporter at the rDNA locus, there is little or no effect of Dsup expression. In contrast, for the *TRP1* reporter at HMR, there is a profound difference between Dsup^−^ and Dsup^+^ cells. While Dsup^−^ grows up to 5 dilutions, Dsup^+^ cells show a strong silencing (∼125-fold difference). We asked if silencing at HMR was associated with a physical change in the chromatin. Inspection of the MNase-seq profile for the HMR region shows a distinct peak at *HMR1* when the nucleosome occupancy is compared between Dsup^−^ and Dsup^+^. This suggests that transcriptional silencing at HMR is correlated with higher nucleosome occupancy in the Dsup^+^ cells and, therefore, lower accessibility for the transcription machinery (Fig. 2c).

**Figure 2.**
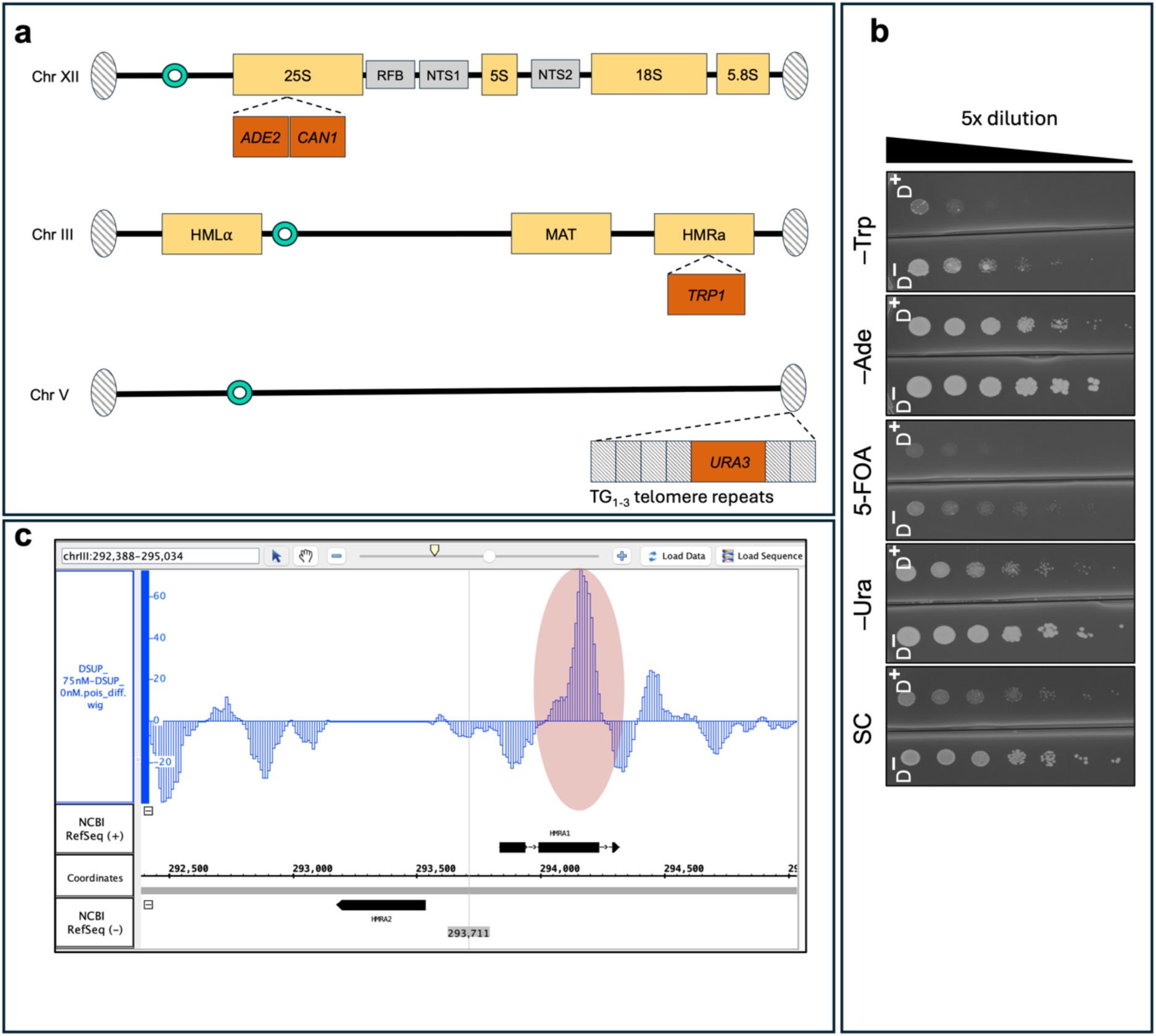
Locus-specific transcriptional silencing loci in yeast. (**a**) In the strain used in this assay (LPY4654), *ADE2–CAN1* reporter is inserted within the 25S rDNA repeat on chromosome XII; RFB denotes the replication fork barrier, and NTS1 and NTS2 are non-transcribed spacer regions. LPY4654 is also MATα, with a *TRP1* reporter integrated at the silent HMRa locus on chromosome III. A third reporter, *URA3*, is positioned adjacent to the right telomere of chromosome V within TG1–3 repeats. When the locus is nucleosome-depleted, silencing is inactive, and therefore transcription can start. Growth on selective media provides a readout of whether silencing at each locus is maintained or relaxed in the presence or absence of Dsup. Hatched ovals indicate telomeres, green circles denote centromeres, and distances and feature sizes in the schematic are not drawn to scale. (**b**) Serial dilution growth assays on complete and selective media. For each plate, D^−^ denotes 0 nM estradiol and D^+^ denotes 75 nM estradiol. (**c**) MNase-seq profile of the HMRa region in strain BY4741 comparing Dsup^−^ and Dsup+ conditions. The y-axis of the plot shows chromatin occupancy. The red oval indicates a region of HMR with an elevated level of chromatin occupancy.

### 2.3 Dsup expression provides higher resistance to oxidative exposure

To examine how Dsup-expressing cells respond to oxidative stress, survival assays were combined with measurements of apoptosis (Fig. 3a). We compared the survival rates of Dsup^−^ and Dsup^+^ cultures following acute hydrogen peroxide exposure. After exposing cultures to 5 mM H_2_O_2_ for 1 h, quantification of survival rates via colony-forming units showed that the Dsup^+^ cultures had a higher survival rate compared to the Dsup^−^ cultures. On average, 139 colonies were observed in the Dsup⁺ samples, compared to only 18 colonies in the Dsup⁻ samples. This corresponds to a ∼8–fold increase in survival in the presence of Dsup protein and demonstrates that Dsup protects yeast cells against oxidative stress.

**Figure 3.**
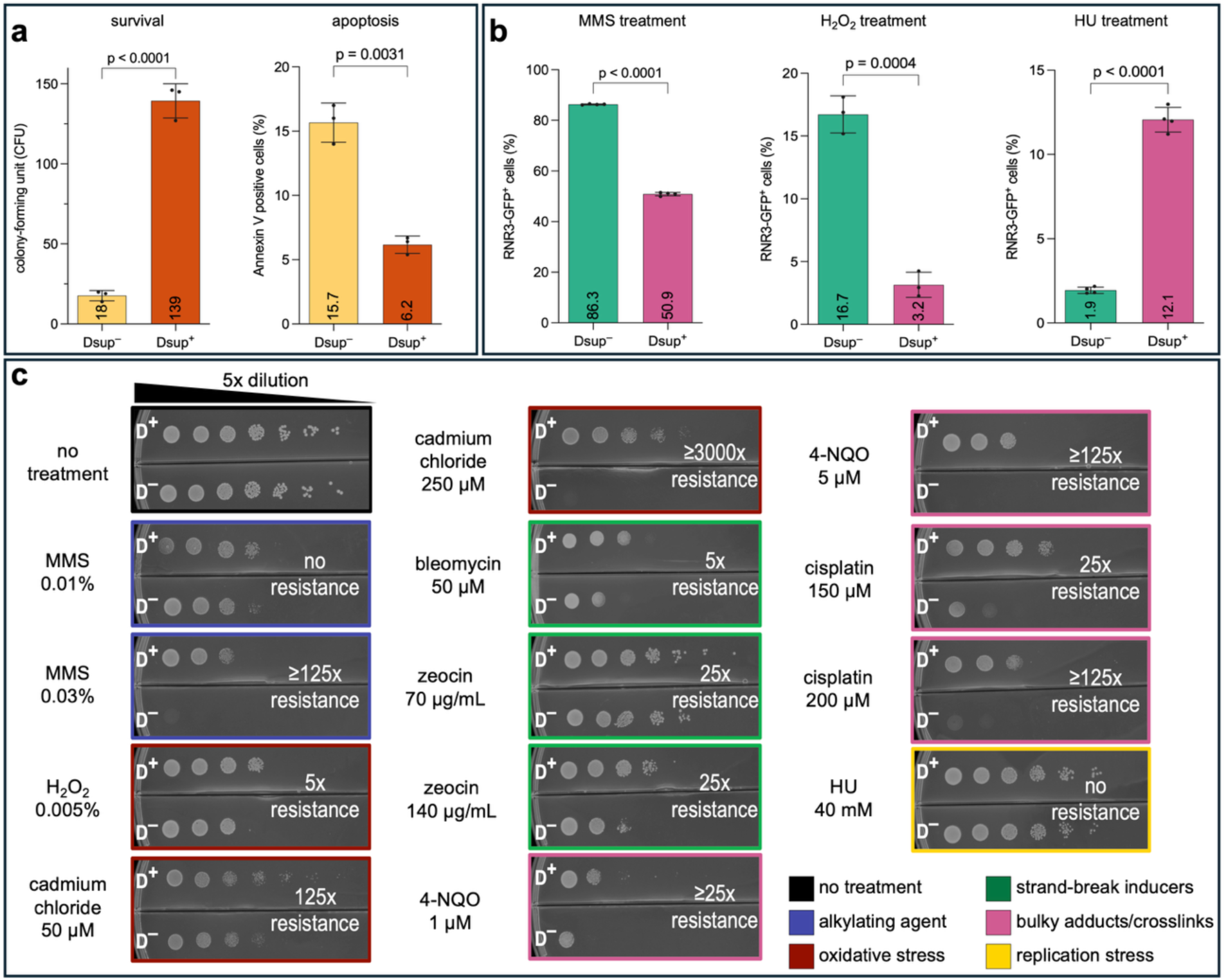
Dsup expression enhances yeast resistance to a variety of DNA-damaging agents. (**a**) Quantification of colony-forming units (CFUs) after treatment with 5mM H2O2; apoptosis rates of cells stained with Annexin-V after treatment with 3mM of H2O2. (**b**) Quantification of Rnr3-positive cells in Dsup⁻ and Dsup⁺ cultures after treatment with MMS (0.015), H2O2 (0.0005%), and HU (15mM-5h). (c) Serial dilutions (5-fold) were prepared starting at OD600 of 0.5 and spotted onto SC in two-compartment Petri dishes containing the indicated concentrations of DNA-damaging agents. Plates with DNA-damaging agents are color-coded by category of DNA damage mechanism.

We then used a lower dose of hydrogen peroxide (3mM) to induce apoptosis, quantified by Annexin V staining. The fraction of apoptotic cells was 15.7% in the Dsup^−^ population and 6.2% in the Dsup^+^ population, a 2.5-fold reduction. Together, these results show that Dsup reduced cell death and also lowered the proportion of apoptotic cells in the population.

### 2.4 Quantification of Rnr3-GFP expression shows Dsup’s bidirectional role under DNA-damaging stresses

Rnr3 expression is a well-established, inducible marker of the DNA damage response in yeast. Because *RNR3* induction is robust and quantifiable, it can serve as a sensitive proxy for DNA damage (*14*). We quantified *RNR3*-GFP induction under three genotoxic conditions to determine if Dsup-mCherry expression alters the DNA damage response in yeast treated with different genotoxins. Treatment with DMSO (2% v/v for 5 and 10 hours) did not activate Rnr3-GFP expression, whereas MMS (0.015%), H_2_O_2_ (0.0005%), or hydroxyurea (15 mM) induced Rnr3 expression. Rnr3-GFP fluorescence was quantified at the earliest detectable time point, *i.e.*, at 5 hours for MMS and HU treatment and at 10 hours for H_2_O_2_.

Under MMS treatment, nearly 86% of Dsup^−^ cells showed GFP signal above background, whereas only 51% of Dsup^+^ cells became GFP-positive, indicating that Rnr3 expression and therefore DNA damage level is lower in Dsup^+^ population. Hydrogen peroxide is primarily an oxidative agent, but prolonged exposure can lead to DNA damage through ROS-mediated base oxidation and strand breaks (*15*). While in the Dsup^−^ population, ∼17% of cells became Rnr3 positive, in the Dsup^+^, the proportion of GFP-positive cells was ∼3%. In contrast to MMS and H_2_O_2_, following treatment with hydroxyurea (HU), *RNR3* induction remained low in Dsup⁻ cells (∼2%) but increased to ∼12% in Dsup⁺ cells, indicating that the presence of Dsup under HU treatment increases DNA damage or replication stress and therefore causes higher checkpoint activation (Fig. 3b).

### 2.5 Dsup Provides Diverse but Not Universal Protection Against DNA Damage

To further assess the impact of Dsup expression on yeast under chronic genotoxic conditions, we performed spot assays with a panel of DNA-damaging agents with diverse mechanisms of action (Fig. 3c). Each spot assay starts with approximately 50,000 cells, followed by serial 5-fold dilutions. In the absence of treatment, Dsup^+^ and Dsup^−^ cultures showed comparable growth. At 0.01% MMS, no substantial growth difference was observed between Dsup^+^ and Dsup^−^ cells. However, at 0.03% MMS, Dsup^+^ maintained growth across several dilutions (25x dilution), whereas Dsup^−^ failed to survive even at the highest inoculum. In oxidative stress conditions (0.005% H_2_O_2_), Dsup^+^ showed 5 times greater survival than Dsup^−^ cells. With 50 μM of cadmium chloride, Dsup^+^ exhibited ∼125-fold higher survival compared to Dsup^−^, and strikingly, with 250 μM cadmium chloride, Dsup^−^ did not grow even at the highest inoculum, while Dsup^+^ continued to grow down to the fifth dilution, indicating at least 3,000 times higher resistance. With the radiomimetic drug bleomycin (at 50 μM), Dsup^+^ showed a 5x higher survival advantage. With zeocin treatment at both 70 μg/mL and 140 μg/mL, Dsup^+^ showed ∼25-fold higher survival compared to the Dsup^−^. With 4-NQO treatment, Dsup^+^ cells showed ≥25x improved resistance at 1 μM. At 5 μM 4-NQO, Dsup^+^ cells were able to form colonies after 3 dilutions (125x dilution) while Dsup^−^ cells were unable to grow even at the highest cell density. With cisplatin treatment, Dsup^+^ retained growth at 150 μM and 200 μM, showing 25- and 125-fold higher survival, respectively, whereas with HU (40 mM), no protective effect was observed, *i.e.*, both Dsup^+^ and Dsup^−^ cells showed comparable growth inhibition.

### 2.6 Chromatin Profiling by MNase-seq

To test the impact of Dsup expression on chromatin structure in yeast, we performed genome-wide MNase-seq in uninduced (0 nM) and induced (75 nM estradiol) conditions. Genome-wide heatmaps of MNase-seq signals of nucleosome occupancy centered at transcription start sites (TSS) (as discussed in references (*16*, *17*)) were used to examine nucleosome occupancy patterns for all genes. Both Dsup^−^ and Dsup^+^ populations displayed the stereotypical phased array of nucleosomes upstream and downstream of the TSS, with no large-scale changes in nucleosome occupancy between the two conditions. Both profiles showed the expected nucleosome-depleted region (NDR) upstream of the +1 nucleosome, a hallmark of active yeast promoters (*18*) (Fig. 4a).

**Figure 4.**
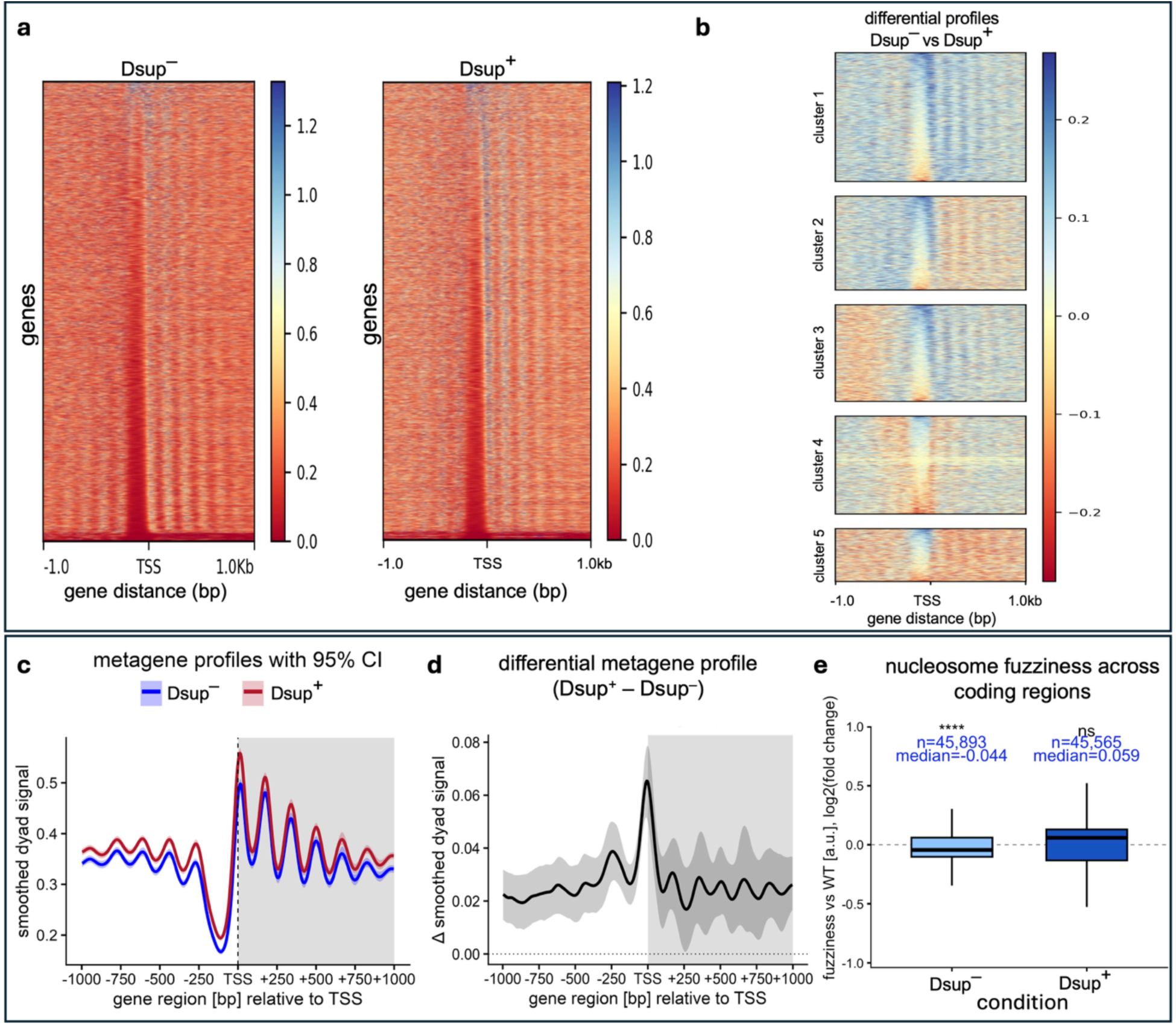
**MNase-seq analysis of nucleosome occupancy and positioning of yeast expressing Dsup**. (**a**) Genome-wide heatmaps of chromatin occupancy aligned to transcription start sites in Dsup^−^ (0 nM estradiol) and Dsup^+^ (75 nM estradiol) conditions. (b) K-means clustering of differential nucleosome occupancy profiles between Dsup⁺ and Dsup⁻ cells. Heatmaps show five distinct clusters of genes based on changes in nucleosome dyad density around TSS ±1 kb. (**c**) Metagene profiles of MNase-seq dyad signal ±1 kb around TSSs for Dsup^−^ and Dsup^+^ samples. The corresponding shaded area represents the 95% confidence interval of the mean. The downstream region of TSS is shaded gray. (**d**) nucleosome occupancy (Dsup^+^ values – Dsup^−^ values) smoothed dyad signal difference across ±1 kb relative to the TSS. The solid line represents the mean difference, while the shaded region indicates the 95% confidence interval derived from bootstrap resampling. The gray background highlights the gene body region (0 to +1000 bp). (**e**) Distribution of nucleosome fuzziness values (log₂ fold-change relative to wild-type) for coding regions passing FDR < 0.05 in Dsup^−^ and Dsup^+^ conditions. Boxplots indicate interquartile range, median, and whiskers (1.5×IQR); “n” denotes the number of nucleosome calls.

To ask if gene expression changes in Dsup^+^ population correlate with chromatin accessibility, we clustered the MNase-seq differential occupancy profiles (Dsup^−^ vs Dsup^+^) using K-means into five groups and then asked if these 5 groups were enriched for certain classes of differentially expressed genes (obtained from RNA-seq). These clusters illustrate distinct patterns of chromatin accessibility around the TSS. Some groups (clusters 1–3) show stronger phasing and prominent nucleosome-depleted regions, while others (clusters 4 and 5) show weaker or more diffuse occupancy signals (Fig. 4b). At the genome-wide scale, we did not observe a correlation between gene expression patterns (|log_2_ fold change| > 1) and occupancy patterns (r = 0.015).

We then used metagene analysis (averaging of signals from all genes) of the dyad signal (*i.e.*, the midpoint of DNA fragments protected by nucleosomes after MNase digestion, which corresponds to the nucleosome center) within ±1 kb of annotated TSSs to compare nucleosome positioning at a population level. Both conditions displayed the expected phased nucleosome arrays downstream of the TSSs. However, Dsup^+^ samples consistently showed a slightly higher dyad signal compared to the uninduced (Dsup^−^) control. The overall phasing and spacing of nucleosomes remained unchanged (Fig. 4c and d). Next, the fuzziness of induced and uninduced samples was compared to the WT control. One-sample Wilcoxon signed-rank test analysis showed a small decrease of fuzziness in Dsup^−^ cells (–0.044 Log_2_FC, p-value <10^-4^), whereas the change in Dsup^+^ population (+0.059 Log_2_FC, p-value = 0.78) is not significantly different from WT samples (Fig. 4e).

### 2.7 Transcriptome analysis shows that Dsup induction rewires metabolism and primes oxidative stress defense

Transcriptome analysis of Dsup^+^ cells (grown in the presence of 75 nM estradiol) revealed substantial changes in gene expression compared to Dsup^−^. A total of 343 and 67 genes were upregulated and downregulated, respectively (adjusted p < 0.01, |log_2_ fold change|>1; Fig. 5a). Inspection of individual candidate genes revealed several functional groups of interest (Fig. 5b). Among the upregulated genes, we noted strong induction of metabolic regulators including *GDH2*, *ACS1*, MPO1 and *GLN1*, as well as nutrient uptake genes with multiple ammonium (*MEP1*/*2*/*3*), amino acid (*PUT1*, *GAP1*, *AGP1*), and sugar transporters (*HXT7*, *HXT9*, *HXT14*) being highly induced. Among the upregulated genes, 18 are involved in the oxidative stress response and localize to different cellular compartments. For example, *GAD1*, *HSP31*, *TSA2*, and *UGA2* are found in the cytoplasm; *FRM2*, *NQM1*, *SRX1*, and *ZTA1* localize to the cytoplasm or nucleus; *DDR2* is present in the cytoplasm or vacuole; *HMX1* resides in the endoplasmic reticulum; and *HYR1*, *TRX2*, and *PRX1* are found in the cytosol, mitochondrial intermembrane space, or mitochondria. While Dsup’s best characterized activity is in protecting chromatin from damage by reactive oxygen species (*5*, *7*), these gene expression changes suggest that even in the absence of external oxidative stress, Dsup activates endogenous redox defense pathways. GO enrichment analysis showed that oxidative stress response, metabolic remodeling, and DNA-related processes are the dominant signatures emerging from the upregulated gene set. Conversely, downregulated transcripts were enriched for genes involved in nucleotide metabolism (*RNR1*, *URA1*/*4*), ribosomal components (*RPL8A*, *RPS26B*, *RPS21A*), and chromatin-related factors (*SPT21*, *HTA2*/*HTB2*). Also, downregulated genes were exclusively enriched for nucleotide and ribonucleoside biosynthetic processes (Table S1). To examine if chromatin accessibility patterns relate to transcriptional activity, we integrated the nucleosome occupancy clusters with RNA-seq expression data. In (Fig. 5c), for each chromatin cluster, genes are represented by their expression fold change (Dsup^−^ vs Dsup^+^), generating transcription heatmaps. To facilitate comparison across genes of different lengths and to focus on transcriptional activity near gene boundaries, we restricted our analysis to a standardized window of ±500 bp around each gene and artificially scaled all ORFs to be the same length to eliminate biases arising from variable gene sizes. This allows consistent visualization of promoter- and terminator-proximal signals and enables detection of antisense or other small RNAs that may accompany expression changes observed within the ORF. Functional enrichment of the genes within each cluster showed that cluster 1 was enriched for pathways linked to metabolism, gene mobility and signaling, regulation of the G2/M transition, and mRNA processing; cluster 2 showed enrichment for genes involved in transcription, while cluster 3 did not show any significant enrichment. Cluster 4 was enriched for pathways related to gene mobility, metabolism, proteolysis, and retrotransposition. Cluster 5 was enriched for carbon metabolism and biosynthesis of secondary metabolites pathways.

**Figure 5.**
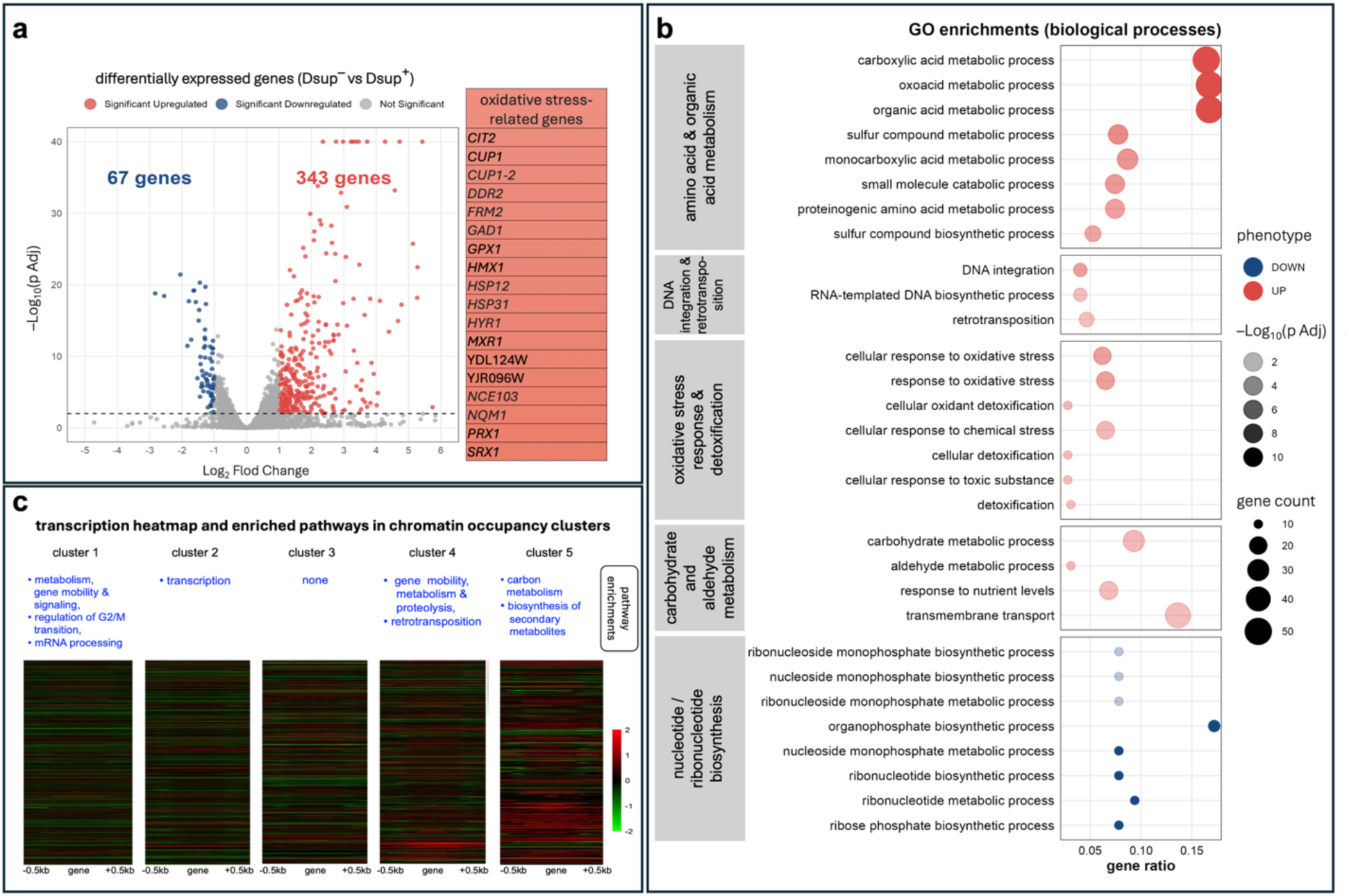
Volcano plot of differential gene expression between Dsup-induced (Dsup⁺) and uninduced (Dsup⁻) cells. (**a**) A total of 343 genes were significantly upregulated (red), and 67 genes were significantly downregulated (blue) at an adjusted p-value < 0.01 and |log₂ fold change| > 1. (**b**) Gene Ontology enrichment of biological processes among differentially expressed genes. (**c**) Transcriptional profiles and functional enrichments corresponding to the five chromatin occupancy clusters. Heatmaps show gene expression changes (Dsup⁺ vs Dsup⁻) for genes within each cluster, where red and green indicate positive and negative log₂ fold changes in expression, respectively.

### 2.8 Synthetic lethality reveals trade-offs of Dsup-mediated DNA protection

To systematically map how Dsup expression affects fitness across diverse cellular processes, we took a Bar-seq screening approach. Specifically, we used the yeast haploid deletion collection engineered via synthetic genetic array (*19*, *20*) to express the Dsup protein. We first identified synthetic lethal interactions, and next, assayed if Dsup modulates mutant fitness in the presence of DNA-damaging stresses, including bleomycin, MMS, and hydrogen peroxide. Such experiments can help generate a genome-wide view of the conditions where the Dsup protein is detrimental or beneficial to the cell. By combining Dsup expression with targeted gene deletions, we can ask which cellular pathways are required for cells to tolerate Dsup. To maintain selection, we grew pools of ∼4300 haploid deletions in SC–HRLK with and without 75 nM estradiol. Significance thresholds were set at 2 for the “Fitness Defect score” in the synthetic lethality screen and for the “Fitness score” in drug treatment assays (fitness score is expressed as the log_2_ difference in the abundance of each strain’s barcode between the test conditions and the controls). 203 gene deletions showed synthetic lethal interactions with Dsup expression at high abundance (Fig 6a). Overall, biological pathways and processes enrichment network reveals several cellular activities: (1) genome-maintenance functions, including DNA repair (BER, post-replication, interstrand cross-link), DNA integrity checkpoints, nucleosome disassembly, negative regulation of DNA replication, and mitotic/meiotic recombination as well as (2) cell-cycle–coupled proteostasis, including APC/C-mediated degradation, ubiquitin recycling, signal-transduction regulation. APC/C-mediated degradation drives key cell-cycle transitions, ubiquitin recycling sustains the proteolytic flux required for those transitions, and signal-transduction pathways modulate checkpoint activation and cell-cycle progression; together linking these processes directly to cell-cycle control.

**Figure 6.**
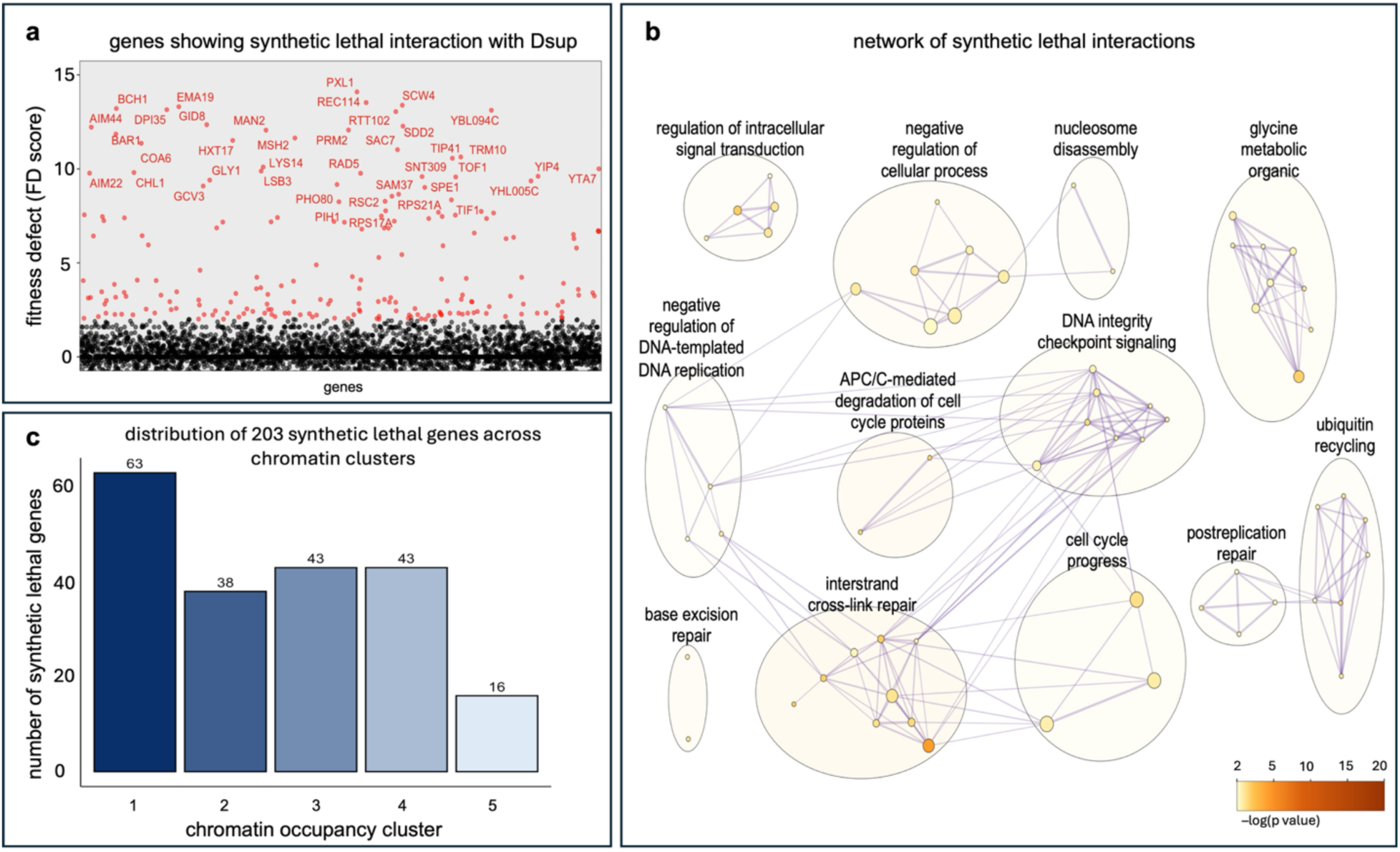
Synthetic lethality profiling of *S. cerevisiae* expressing Dsup. (**a**) Scatterplot of gene deletion strains showing fitness defects (y-axis, FD score) upon Dsup induction (75 nM estradiol) compared to uninduced controls. With a threshold of 2 units of log2 foldchange used to define significant growth difference (shown in red), higher values indicate greater sensitivity, *i.e.*, stronger synthetic lethal interactions with Dsup. The top 40-scoring deletions are labeled in each plot. (**b**) Network representation of enriched pathways from Dsup synthetic lethal interactions. Genes identified as synthetic lethal with Dsup expression were subjected to pathway and process enrichment analysis using Metascape (*20*). The plot shows representative GO Biological Process and pathway terms (p < 0.01, enrichment factor > 1.5), clustered by membership similarity (edges represent kappa score > 0.3). Each node denotes an enriched term. (**c**) The bar plot shows how many of synthetic lethal genes belong to each nucleosome occupancy profile.

To determine if there is a relationship between these synthetic lethal gene deletions and their chromatin occupancy (*i.e.* do sensitive gene deletions display altered chromatin architecture in wild type cells expressing Dsup), we investigated differential occupancy profiles of them (Fig. 4b). Of 203 genes, 144 synthetic lethal deletions (71%) belong to clusters 1, 2, and 3, which have strong (fixed) phasing and prominent nucleosome-depleted regions. 59 synthetic lethal deletions are from clusters 4 and 5 that show weaker or more diffuse occupancy signals (Fisher’s exact test, p = 0.009). This indicates that genes that are essential for overcoming the stress caused by Dsup tend to have higher nucleosome occupancy when they exist in WT (Fig. 6c) (Table S2). Synthetic lethal genes exhibited a protein abundance distribution that closely mirrored that of the global proteome reported on PaxDb (*21*). A chi-square goodness-of-fit test comparing abundance-class frequencies revealed no substantial deviation across most abundance ranges, indicating that synthetic lethality is not broadly associated with protein abundance. Minor deviations observed in the highest-abundance classes were not considered further due to low expected counts.

### 2.9 Dsup enhances yeast survival under multiple genotoxic stresses through distinct pathway-specific mechanisms

For Bar-seq screens under DNA-damaging conditions, we induced Dsup expression at 40 nM β-estradiol rather than 75 nM to obtain a more modest Dsup-associated fitness defect and increase the dynamic range of the drug assays.

#### Bleomycin Treatment

The bleomycin Bar-seq screen was performed for 5 generations of growth. The results show that Dsup expression enhances yeast survival in the presence of bleomycin-induced stress, which is known to induce ROSs as well as single-or double-stranded DNA breaks (*22*). Deletions that benefited from Dsup expression were enriched for GO terms that are known to be activated under bleomycin-induced stress in yeast and mammalian cells (Table 1). These include vesicle-mediated transport (*vps1*τι, *vps24*τι, *vps8*τι, *ypt6*τι) (*23*, *24*), vacuolar trafficking (*vam10*τι, *prb1*τι, *vps35*τι) (*23*, *24*), membrane organization (*gyp1*τι, *rvs167*τι, *gvp36*τι), which is an indirect consequence of bleomycin treatment (*25*), macroautophagy (*TRS85*τι, *VPS38*τι) (*26*), and autophagy (*prb1*τι, *cue5*τι, *doa1*τι, *sac1*τι, *sic1*τι, *trs85*τι) (*26*, *27*). Another enriched cellular component GO term is endomembrane system, which provides the membrane sources and trafficking machinery for autophagy and vacuolar pathways (*28*, *29*). We also observed ESCRT, Golgi, and vacuolar enrichments. Each of these processes is involved in sorting proteins for degradation.

**Table 1.**
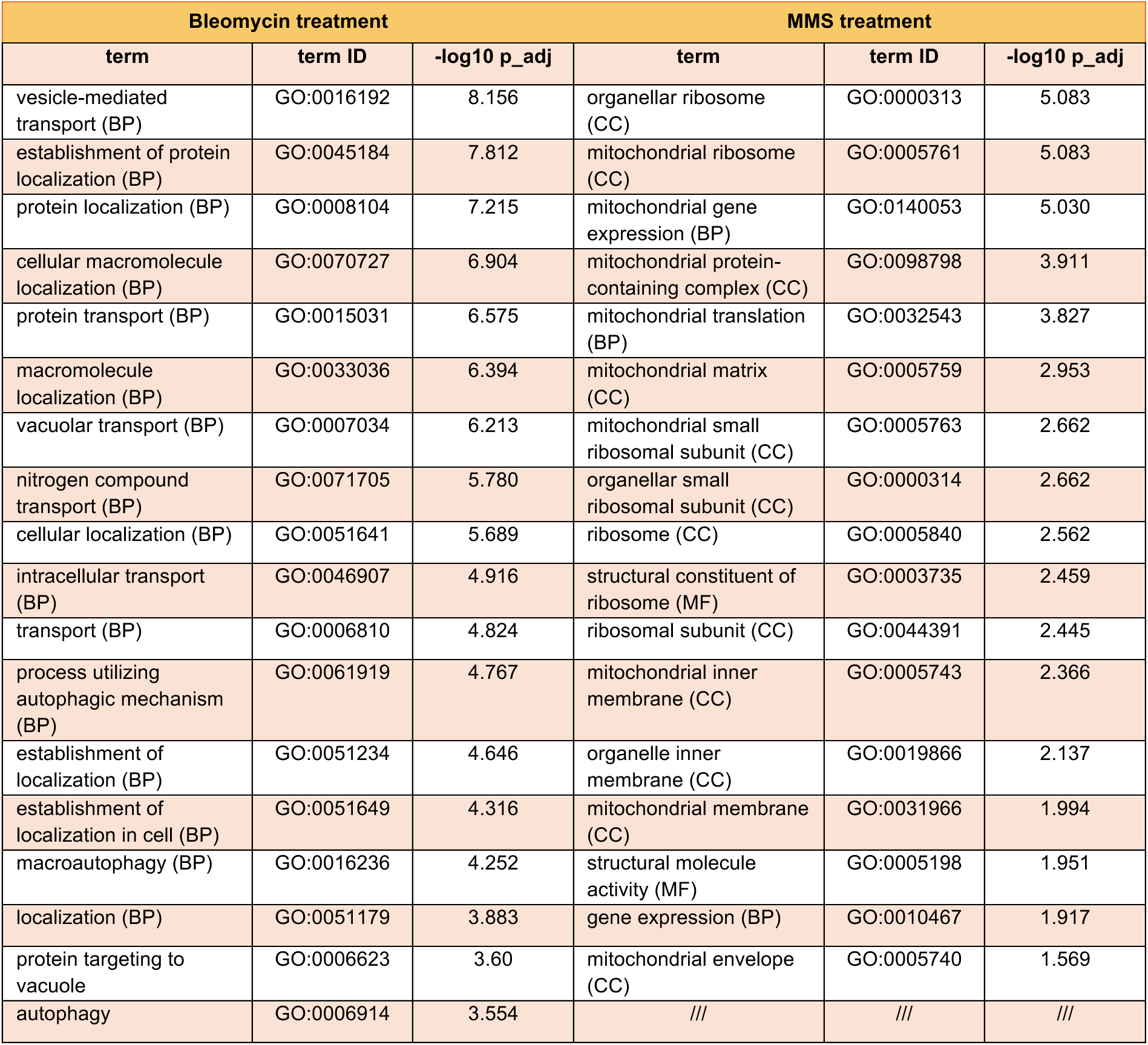
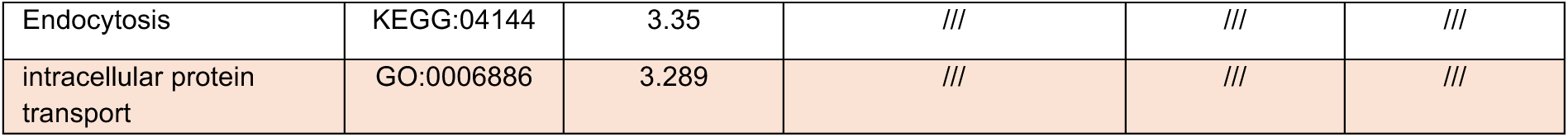
Top 20 GO enrichment of bleomycin and all GO enrichments of MMS treatment for deletions whose fitness improved in the presence of Dsup protein. .

| Bleomycin treatment |  |  | MMS treatment |  |  |
| --- | --- | --- | --- | --- | --- |
| term | term ID | -log10 p_adj | term | term ID | -log10 p_adj |
| vesicle-mediated transport (BP) | GO:0016192 | 8.156 | organellar ribosome (CC) | GO:0000313 | 5.083 |
| establishment of protein localization (BP) | GO:0045184 | 7.812 | mitochondrial ribosome (CC) | GO:0005761 | 5.083 |
| protein localization (BP) | GO:0008104 | 7.215 | mitochondrial gene expression (BP) | GO:0140053 | 5.030 |
| cellular macromolecule localization (BP) | GO:0070727 | 6.904 | mitochondrial protein-containing complex (CC) | GO:0098798 | 3.911 |
| protein transport (BP) | GO:0015031 | 6.575 | mitochondrial translation (BP) | GO:0032543 | 3.827 |
| macromolecule localization (BP) | GO:0033036 | 6.394 | mitochondrial matrix (CC) | GO:0005759 | 2.953 |
| vacuolar transport (BP) | GO:0007034 | 6.213 | mitochondrial small ribosomal subunit (CC) | GO:0005763 | 2.662 |
| nitrogen compound transport (BP) | GO:0071705 | 5.780 | organellar small ribosomal subunit (CC) | GO:0000314 | 2.662 |
| cellular localization (BP) | GO:0051641 | 5.689 | ribosome (CC) | GO:0005840 | 2.562 |
| intracellular transport (BP) | GO:0046907 | 4.916 | structural constituent of ribosome (MF) | GO:0003735 | 2.459 |
| transport (BP) | GO:0006810 | 4.824 | ribosomal subunit (CC) | GO:0044391 | 2.445 |
| process utilizing autophagic mechanism (BP) | GO:0061919 | 4.767 | mitochondrial inner membrane (CC) | GO:0005743 | 2.366 |
| establishment of localization (BP) | GO:0051234 | 4.646 | organelle inner membrane (CC) | GO:0019866 | 2.137 |
| establishment of localization in cell (BP) | GO:0051649 | 4.316 | mitochondrial membrane (CC) | GO:0031966 | 1.994 |
| macroautophagy (BP) | GO:0016236 | 4.252 | structural molecule activity (MF) | GO:0005198 | 1.951 |
| localization (BP) | GO:0051179 | 3.883 | gene expression (BP) | GO:0010467 | 1.917 |
| protein targeting to vacuole | GO:0006623 | 3.60 | mitochondrial envelope (CC) | GO:0005740 | 1.569 |
| autophagy | GO:0006914 | 3.554 | /// | /// | /// |
| Endocytosis | KEGG:04144 | 3.35 | /// | /// | /// |
| intracellular protein transport | GO:0006886 | 3.289 | /// | /// | /// |

Pathway- and process-level enrichment showed a network of terms that are directly related to autophagy (vesicle-mediated transport; process utilizing autophagic mechanism; vesicle organization; endosomal transport; membrane trafficking; endomembrane system organization). Additionally, processes that indirectly contribute to autophagy (protein localization to the Golgi apparatus; intra-Golgi vesicle-mediated transport; establishment of protein localization to membranes) (Fig. 7) are enriched. These processes are indirectly related to autophagy because the Golgi and associated trafficking pathways supply membranes, lipids, and sorting machinery required for autophagosome formation and maturation. Proper localization of proteins to the Golgi or other membranes ensures that ATG proteins, vesicle-fusion factors, and lipid-modifying enzymes reach the compartments that support autophagic flux (*30*, *31*) (Table S3).

**Figure 7.**
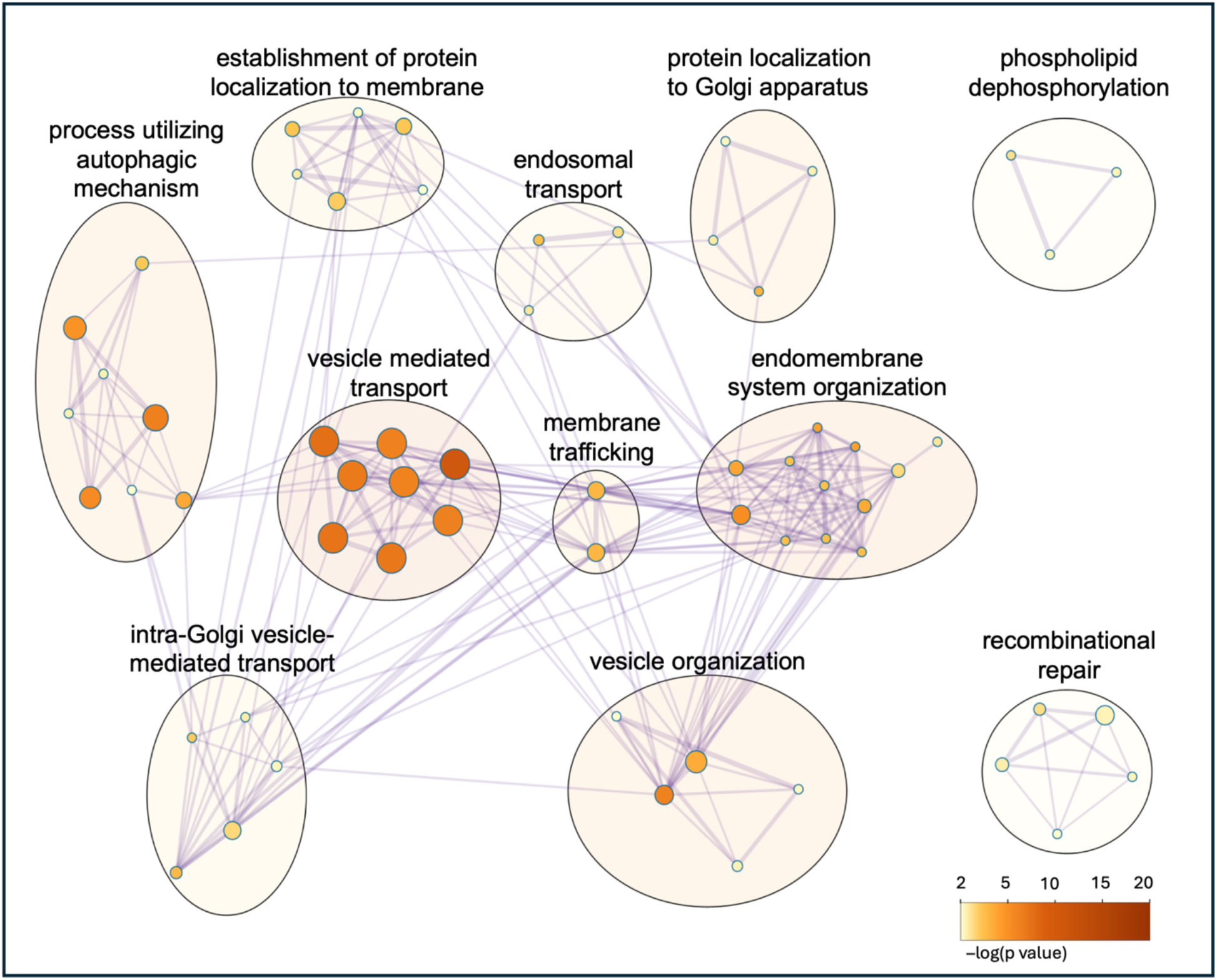
Process and pathway enrichment for mutants whose fitness was enhanced by Dsup during exposure to bleomycin. Enrichment was performed using Metascape (p < 0.01, enrichment factor > 1.5). Nodes represent enriched terms, clustered by membership similarity (edges represent kappa score > 0.3).

#### MMS treatment

Expression of Dsup in *S. cerevisiae* afforded protection against MMS-mediated DNA damage (Fig. 3b). It has been suggested that DSBs arise after MMS treatment when, during BER, single-strand breaks (SSBs) are encountered by a replication fork (*32*). The Bar-seq screen with MMS showed that Dsup induction increased the fitness of several deletion strains, notably *rad2*τι, *pbr1*τι, and *htz1*τι strains. GO enrichment analysis for strains with improved fitness (fitness score >2) revealed significant overrepresentation for mitochondria-related terms (mitochondrial gene expression; mitochondrial translation; mitochondrial ribosome; mitochondrial matrix; mitochondrial membrane; mitochondrial envelope) and ribosomal structure-related terms such as structural constituent of ribosome, organellar ribosome, mitochondrial ribosome, mitochondrial small ribosomal subunit, organellar small ribosomal subunit, as well as ribosomal subunit but not direct DNA repair terms (Table 1). We also observed enrichment for chromatin-related gene deletions such as *htz1*τι (variant H2A.Z) and *hht2*τι (histone H3 in yeast), both of which are known to dictate transcriptional outcomes and DNA accessibility upon stress (*33*, *34*) (Table S4). Table 1 shows the GO enrichments of deletion mutants benefiting from Dsup expression under the same treatment. Our data reveal that Dsup reduces MMS-induced DNA damage in yeast by reducing reliance upon NER mechanism (*RAD2* and *HTZ1*) and stabilizing ribosomal and mitochondrial functions (*35*).

#### Hydrogen peroxide treatment

We identified 55 mutants whose growth was enhanced during oxidative (hydrogen peroxide) stress in the Dsup^+^ deletion pool compared to the uninduced pool (Table S5). These mutants represent a broad set of cellular activities. Among these were gene deletions involved in cell cycle and genome maintenance, such as *ace2*Δ, *ctf4*Δ, and *mrc1*Δ; chromatin and transcription regulation, such as *arp6*Δ, *sin3*Δ, and *src1*Δ; mitochondrial/metabolic activities, such as *cox10*Δ, *cox18*Δ, *cyt1*Δ, *mgr1*Δ, *shy1*Δ, and *pos5*Δ. Other groups represented were transport/trafficking proteins (*alp1*Δ, *fen2*Δ, *fps1*Δ, *gpr1*Δ, *gyp1*Δ, *ist1*Δ, *itr1*Δ, *hut1*Δ, *tvp18*Δ), ribosomal proteins (*rpl13b*Δ, *rps18b*Δ, *rps30a*Δ, *rtc6*Δ), and RNA processing deletions (*cdc40*Δ, *lsm6*Δ, *sac3*Δ). Some enzyme and protein modifier deletions like *apa2*Δ, *eug1*Δ, *hrf1*Δ, *pfa4*Δ, *prb1*Δ, *spr6*Δ, and *yke2*Δ were also among those showing gained fitness in the Dsup^+^ pool. Notably, *EUG1* is an ER disulfide isomerase, and its deletion has been associated with reduced replicative lifespan, a phenotype commonly associated with impaired proteostasis and redox homeostasis in oxidative stress–associated replicative aging (*36*). Even though certain functional tendencies were observed (*e.g.*, replication, chromatin, nuclear organization), formal GO enrichment analysis did not identify any statistically significant categories. Specifically, while several terms showed p-values corresponding to -log_10_ (p) ≈ 2–3, none surpassed the multiple-testing correction threshold (q < 0.05), which is the standard cut-off for statistical significance. As a result, further interpretation was focused on gene-level patterns, rather than pathway-level enrichment in the discussion.

## 3. Discussion

In this study, we investigated the effects of heterologous expression of the tardigrade protein Dsup in *S. cerevisiae* using growth assays, DNA damage reporters, chromatin profiling, transcriptomics, and genome-wide genetic interaction screens. We found that Dsup broadly enhances resistance to genotoxic and oxidative stress, while modestly perturbing chromatin accessibility, slowing cell-cycle progression, and creating specific fitness costs and genetic dependencies.

One possible explanation for the rapid gene silencing of Dsup on low-copy ARS/CEN6 plasmids with strong promoters (constitutive *TDH3* or inducible *GAL1*) is that cells actively repress Dsup through transcriptional or post-transcriptional mechanisms. This would be consistent with the fact that high-level Dsup expression makes cells quite sick. This phenomenon has been reported for other heterologous proteins expressed in yeast (*37*). Galactose-induced expression of Dsup resulted in a prolonged lag phase, and when logarithmic growth commenced, Dsup expression was largely silenced, with Dsup detectable in only ∼2-3% of cells. A recent study by Fujita *et al.* showed that strong protein expression in yeast leads to reduced growth rate, and even non-toxic proteins can impose a substantial translational and metabolic burden when expressed at high levels (*38*). Dsup’s effects seem to go beyond a generic proteotoxic burden, especially when compared to the GFP control, which showed a more modest growth defect with no effect on the lag phase. These findings showed that high-level Dsup expression was very unstable and motivated us to switch to a tunable expression strategy. After observing a heterogeneous expression from a centromeric plasmid with an estradiol promoter, we integrated the construct into the genome, which yielded a more durable and homogenous expression. With 75 nM of estradiol induction, expression reached a relatively high level while causing a moderate fitness defect of about 14% (Fig. 1i). Even with this well-tolerated expression strategy, Dsup^+^ cells accumulate in G2/M phase in a dose-dependent manner. This agrees with the earlier observation that induction of *GAL1* promoter caused a prolonged G2/M. The consistent G2/M arrest suggests that Dsup slows DNA replication or triggers checkpoints.

Our results show that Dsup expression in yeast protects against DNA-damaging agents with a variety of mechanisms. In survival assays and oxidative stress-induced apoptosis, Dsup proved to be protective, and cells equipped with Dsup demonstrated both higher survival and a lower rate of programmed cell death (Fig. 3a). By reducing DNA lesions, Dsup likely prevents the DNA damage signals that trigger apoptosis. In spot assays, Dsup^+^ cells survived far better when exposed to MMS, hydrogen peroxide, bleomycin, 4-NQO, cadmium, zeocin, and cisplatin (Fig. 3c). This broad protection is a novel observation that expands beyond the oxidative and radiation stress protection previously reported (*5*, *39*). However, Dsup conferred no survival benefit under HU treatment. Moreover, the observation with *RNR3*-GFP reporter showed that under HU treatment, the presence of Dsup increases the DNA damage and therefore checkpoint activation. HU is known for primarily stalling replication forks by depleting dNTP pools, but it also causes DNA double-strand breaks and accumulation of recombination intermediates (*40*). A recent study on yeast showed that HU inhibits DNA polymerase complexes by oxidizing iron-sulfur (Fe-S) clusters, rendering them unable to bind DNA (*41*). This fits a model where Dsup binds DNA and, by acting as a physical barrier, prevents damage from ROSs or other DNA-damaging agents. In HU treatment, however, Dsup’s presence appears to further obstruct replication. At the same time, the lower rate of Rnr3 expression under MMS and H_2_O_2_ treatment suggests that fewer lesions occur in cells when Dsup is present, as Dsup physically protects cells from alkylation and free-radical attack (Fig. 3b). The effect of Dsup on heterochromatin in a locus-specific manner implies that Dsup does not globally alter all heterochromatin (Fig. 2b). Dsup contains an HMGN-like nucleosome-binding domain (*7*), so it has a mechanism to locally alter chromatin. In support of this, Dsup expression has been reported to promote chromatin condensation in mammalian neurons (*11*). It is plausible that Dsup coating of DNA further represses chromatin at HMR and enhances silencing at that locus. This result should be interpreted with caution, because the MNase-seq was performed in BY4741 (*MATa*), whereas the silencing assay was carried out in MATα (LPY4654 (*13*)). However, since rDNA and telomeric markers show that silencing at these loci was unchanged, we can infer that Dsup’s effect on chromatin silencing is not universal. Instead, Dsup might compete with transcription factors or chromatin remodelers at specific sites, as seen in other models, such as *D. melanogaster* (*2*, *10*).

In MNase-seq analysis of Dsup^−^ vs Dsup^+^ conditions, chromatin occupancy heatmaps, metagene profiles, and fuzziness distributions reveal subtle differences in nucleosome positioning and occupancy (Fig. 4). Also, the experiments with silencing markers (such as the HMR locus) suggest that such changes in occupancy may be localized rather than global remodeling. Clustering analysis of differential occupancy profiles showed that genes in Cluster 5 (characterized by reduced nucleosome occupancy around TSSs) showed a stronger overlap with genes that are upregulated by Dsup expression (Fig. 4b). 71% of the synthetic lethal gene deletions belonged to clusters 1–3, which display stronger phasing and permanent nucleosome-depleted regions. Also, cluster 1 was enriched for ‘regulation of the G2/M transition’, the stage of cell cycle that is affected by Dsup expression under galactose-induced expression. Together, these observations suggest that genes whose deletion causes synthetic lethality with Dsup (that is, genes required for cells to tolerate Dsup expression) may preferentially exhibit strong chromatin phasing and more stable nucleosome-depleted regions.

In the metagene analysis, there was a consistent enhancement of the dyad signal in Dsup^+^ (Fig. 4c and d), and the fact that Dsup physically coats the chromatinized genome suggests that Dsup association may enhance protection of DNA from MNase digestion. In fuzziness analysis, although compared to WT cells, Dsup^−^ cells showed a reduced fuzziness (Fig. 4d), this alteration is minor and perhaps negligible, which might be due to experimental noise or subtle chromatin fluctuations unrelated to Dsup expression.

In the RNA-seq experiment, Dsup expression led to an upregulation of genes involved in the cellular response to oxidative stress, including both general and organelle-specific detoxification pathways, which suggests that Dsup expression appears to activate endogenous redox defense systems (Fig. 5b). If true, this observation may reflect a form of molecular preconditioning (*42–45*), in which the presence of Dsup primes the cell’s antioxidant capacity and enhances its resilience against subsequent stresses. Dsup may not only shield DNA but also boost cellular stress responses through gene regulation. Upregulation of core metabolic processes in Dsup^+^ (including carboxylic acid metabolic process, oxoacid metabolic process, and organic acid metabolic processes) shows a broad-based increase in turnover of organic acids. This pattern could reflect the cell’s attempt to meet the higher energy and biosynthetic burden associated with high-level Dsup expression. Moreover, upregulation of sulfur compound metabolic process suggests activation of glutathione-dependent detoxifying machinery (*e.g.*, upregulation of *GTT1*, *TRX2*, and *GPX1*) or related sulfur metabolism pathways for antioxidant defense. Together, these signatures suggest Dsup^+^ cells enter a metabolically challenging and stress-prepared state even in the absence of exogenous DNA damage.

Conversely, downregulation of nucleotide synthesis pathways in Dsup+ cells likely reflects slower proliferation under 75 nM estradiol induction, reducing demand for DNA and RNA synthesis. This shift may indicate a reallocation of resources from nucleotide production toward stress responses, protein maintenance, and metabolic adaptation, consistent with a slower-growing but more stress-resilient physiological state. For instance, Lu *et al.* reported that slower growth induced heat-shock resistance in yeast (*46*), and Zakrzewska *et al.* reported that “slow growth, whether induced by mutation during exponential growth or induced by nutrient-limited growth in chemostats, led to an increased tolerance for oxidative, acid, or heat stress” (*47*).

The synthetic lethal screen revealed that expressing Dsup in yeast causes specific vulnerabilities, which manifest differently when combined with different deletion strains. Deletions of DNA repair genes, chromatin remodelers, and transcription regulators became selectively lethal when Dsup was expressed at a high level. Dsup can prime cells to cope with stress but is equally adept at altering pivotal genome functions. The picture that emerges is a trade-off. Protection is gained, but the cell now relies more heavily on certain pathways to keep growing. The strongest dependency is DNA Damage Response and DNA repair—*rad1*Δ*, rad5*Δ*, rad9*Δ*, rad17*Δ*, rad18*Δ*, rad51*Δ, and corresponding gene deletions are permissible in normal situations but lethal with Dsup. A simple reading is that Dsup decreases the number of breaks, but access to the remaining lesion is perhaps made more restricted. Any DNA break (especially double-strand breaks) that slips through now requires high-fidelity repair; without it, instability is enhanced, and the cell perishes. Analogous outcomes have been observed in neurons, where Dsup can enhance damage levels rather than reduce them (*11*).

Deletion of chromatin remodelers *INO80* and *ARP8* also showed synthetic lethality with Dsup expression. The Ino80 chromatin-remodeling complex shifts nucleosomes during transcription and DNA repair, and its Arp8 subunit facilitates access to the complex and DNA (*48–50*), indicating that Dsup, when there is insufficient remodeling activity, can render cells more susceptible to replication stress.

Also, deletion of *MED1* showed synthetic lethality with Dsup expression, which is a transcription activation mediator subunit for promoting initiation by RNA polymerase II and helping with stress-related gene expression regulation (*51*). This observation suggests that Dsup can interfere with gene expression, perhaps by repressing promoter access or stalling polymerases. This is consistent with another report in *D. melanogaster* on Dsup’s as a general transcriptional repressor (*10*)

Bleomycin generates ROS and induces single- and double-stranded DNA breaks(*22*). In bleomycin Bar-seq screens, deletions that showed enhanced fitness upon Dsup expression were enriched for vesicle-mediated transport, vacuolar transport, autophagy-related processes, the endomembrane system, and ESCRT components. Recombinational repair factors were also enriched, consistent with reports that autophagy can act upstream of recombination by clearing DNA-blocking protein–DNA lesions (*52*). These results support a scenario in which Dsup reduces the initial damage burden, which in turn decreases reliance on autophagy and recombination pathways in bleomycin stress. Disruption of autophagy or vacuolar transport may further enhance Dsup’s protective effects by increasing its stability or persistence, as these pathways can contribute to the degradation or sequestration of exogenous proteins via bulk autophagy (*53*).

In Bar-seq under MMS treatment, deletions with enhanced fitness due to Dsup expression showed overrepresentation for a number of mitochondria-related and ribosomal structure-related GO terms (Table 1). As dysfunctional mitochondria further increase oxidative stress, Dsup’s protection to the genome may indirectly help by inhibiting formation of second ROS generation. This result is consistent with our other observations using the *RNR3*-GFP DNA damage marker, which showed that Dsup^+^ cells exhibit reduced MMS-induced activation of *RNR3*. These findings confirm Dsup’s potential double protection mechanisms: (i) direct protection of chromatin by nucleosome binding (*5*), and (ii) indirect regulation of cellular stress responses by antioxidant defenses (*8*), and chromatin reorganization (*12*) through preserving genome integrity and enhancing stress resistance. The result of Bar-seq under H_2_O_2_ treatment suggests that Dsup’s benefit during oxidative stress affects multiple pathways. Replication and checkpoint-deficient strains (such as those with deleted *CTF4* and *MRC1*) that are hypersensitive to DNA damage showed enhanced fitness when Dsup was expressed. Here, Dsup’s protection would reduce accruing DNA lesions and thus provide partial compensation for the checkpoint and replication defects. Similarly, mutants with loss of function of chromatin remodelers and transcriptional regulators (*arp6*τι*, sin3*τι, and *src1*τι), whose fitness was improved with Dsup expression, may tolerate increased genomic instability. In other words, Dsup binding to nucleosomes could help stabilize DNA in these backgrounds, though it would not restore the lost regulatory activities. For mitochondrial and metabolic deletions, such as *cox10*τι*, cyt1*τι, and *pos5*τι, Dsup’s benefit is likely to be indirect. Perturbation of these activities imposes enhanced oxidative stress through dysfunctional respiration and/or NADPH delivery. Dsup alone cannot rescue metabolic capacity but can prevent the consequent damage to nuclear DNA integrity and thus explain some viability improvement of those deletions.

From these observations, we propose a model whereby Dsup wraps around chromatin to shield DNA and reduce oxidative and genotoxic stress damage. This interaction with chromatin could limit access of accessory proteins, slowing replication forks, altering transcription, and reducing DNA repair. Interestingly, the HU Bar-seq screen by Lee *et al.* shows enrichments for DNA repair processes (such as DNA repair, recombinational repair, and double-strand break repair) (*54*). We infer that the toxicity of HU and Dsup overlaps in specific DNA repair pathways, which explains why Dsup, combined with HU treatment, caused higher induction of *RNR3* (Fig. 3b).

This model makes testable predictions. For instance, replication forks in Dsup^+^ cells will progress slowly. Therefore, a Dsup mutant lacking the nucleosome-binding motif (HMGN-like nucleosome-binding sequence identified by Chavez *et al.* (*7*)) should lose its protective function and also not show synthetic lethality with as many DNA repair/replication processes anymore.

The phenotypes we observe in yeast are mainly consistent with other models. Mammalian HEK293 cells expressing Dsup showed increased X-ray and H_2_O_2_ tolerance, and reduced DNA damage (*5*). In *D. melanogaster*, although Dsup expression caused higher survival under oxidative stress and longer lifespan, flies had reduced locomotor activity and extensive downregulated expression of neural genes (*10*). This indicates some stress or toxicity at high Dsup levels. Notably, rat neurons expressing Dsup showed the opposite effect—increased DNA breaks, chromatin condensation, and cell death (*11*). Finally, plant cells expressing Dsup showed reduced UV-induced DNA damage (*55*).

Across yeast and animal models, Dsup consistently acts as a chromatin-associated DNA shield, but the extent and consequences of this protection diverge. In yeast, both our study and Aguilar *et al.* agree that Dsup enhances survival under oxidative stress. However, they interpret this effect as narrow and largely cost-free (reduced fitness), attributing protection exclusively to DNA shielding, with no transcriptional priming, ROS scavenging, or growth defects (*12*). In contrast, our yeast study reveals a broader role. Dsup confers protection against diverse genotoxins, induces oxidative stress–response processes even in the absence of stress, slows growth, delays the cell cycle, and causes dependencies on DNA repair-related pathways. Extending beyond yeast, the study by Richaud *et al.* demonstrates that Dsup expression in *C. elegans* delays ROS accumulation via reduced mitochondrial respiration, increases resistance to oxidative stress and ionizing radiation, and extends lifespan. They also report no overt fitness costs at the organismal level. Without transcriptomic analyses, however (*9*); their conclusions are limited to specific signaling and physiological observations.

In evolutionary terms, our results, along with previously published data, suggest that tardigrade Dsup did not evolve in isolation, but rather as a response to its occasionally extreme lifestyle and environment. For example, the fitness benefits and costs of robust DNA repair, chromatin-modulating, and transcriptional regulation activities would all have to be taken into consideration. In yeast, where those co-adaptations are absent, Dsup expression reveals hidden fragilities in both wild type and mutant genomes. This study provides a resource describing these diverse “Dsup dependencies” that must be accounted for to realize the biotechnological or therapeutic benefits of transgenic Dsup. For medicine and biotechnology applications, the lesson is caution. Dsup can boost defense to DNA damage but introduce new dependencies or susceptibilities unless accompanied by compensation.

## 4. Methods

Strain and plasmids used in this study are listed in Table 2.

**Table 2.** Strains and plasmid used in this chapter.

| strain / plasmid | genotype / features |
| --- | --- |
| BY4741 | *MATa his3Δ1 leu2Δ0 met15Δ0 ura3Δ0* |
| yGWB3851 | *MATa RNR3::GFP[HIS3MX] leu2Δ0 his3Δ1 ura3Δ0 met15Δ0* |
| LPY4654 | *MATα ade2-1 can1-100 his3-11 leu2-3,112 trp1-1 ura3-1 GAL rDNA::ADE2CAN1 hmr:: TRP1 VRTEL::URA3* |
| HKG_Sc_046 | *MATa COX4::mCherry[KanMX] leu2Δ0 his3Δ1 ura3Δ0 met15Δ0* |
| Y7092 | *MATα can1Δ::STE2pr-Sp_his5 lyp1Δ ura3Δ0 leu2Δ0 his3Δ1 met15Δ0* |
| HKG_pSc_07 | p*TDH3*, Dsup-EGFP, tADH1, *LEU2* selection, CEN6; E.coli_AmpR |
| HKG_pSc_28 | p*TDH3*, EGFP, tADH1, *LEU2* selection, CEN6; E.coli_AmpR |
| HKG_pSc_22 | p*GAL1*, Dsup-EGFP, tADH1, *LEU2* selection, CEN6; E.coli_KanR |
| HKG_pSc_27 | p*GAL1*, EGFP, tADH1, *LEU2* selection, CEN6; E.coli_KanR |
| HKG_pSc_103 | pEstradiol, Dsup-EGFP, tADH1, integrates *LEU2* marker at *LEU2* locus; E.coli_KanR |
| HKG_pSc_108 | pEstradiol, Dsup-mCherry, tADH1, integrates *LEU2* marker at *LEU2* locus; E.coli_KanR |

### Media

Experiments involving centromeric plasmids were performed in synthetic complete medium lacking leucine (SC–Leu). For integrating plasmids, transformants were initially selected on SC–Leu plates; following confirmation of genomic integration, all subsequent characterizations were performed in complete SC medium without selection. For Bar-seq experiments, pools were grown in SC–HRLK+ canavanine / thialysine /G418 with either 75 nM (in synthetic lethality screen) or 40 nM (for drug treatments) of estradiol.

### Cloning, yeast transformation, and integration confirmation by colony PCR

The Dsup sequence was codon-optimized for *S. cerevisiae* using IDT Codon Optimization Tool and synthesized by IDT with overhangs for Golden Gate assembly. Plasmids were assembled de novo using modules from the Yeast MoClo toolkit together with a supplementary module kit providing an estradiol-inducible expression system (*56*, *57*). Reaction designs were generated using the NEB Golden Gate Assembly Tool (v2.7.1). DNA assemblies were performed following the NEB Golden Gate (24-fragment) Assembly Protocol. Briefly, equimolar concentrations of DNA modules were mixed in a 25 µL reaction containing 1X T4 DNA ligase buffer, 0.5 µM T4 DNA ligase (NEB #M0202), and 1.5 U BsaI-HFv2 (NEB #R3733). Assembly reactions were cycled in a thermocycler under the following program: (37 °C for 5 min → 16 °C for 5 min) × 30 cycles, followed by 60 °C for 5 min.

Following Golden Gate assembly, 10 µL of the reaction mix was transformed into *E. coli* (DH5α) and plated on LB agar containing either ampicillin or kanamycin. Plasmids were sequence-verified (Plasmidsaurus) and subsequently used for yeast transformation. For integration constructs, plasmids were linearized with NruI (NEB #R3192S) prior to yeast transformation. Yeast competent cells were prepared using the Frozen-EZ Yeast Transformation II Kit (Zymo Research T2001), following the manufacturer’s standard protocol. Chromosomal integration of transformation cassettes at the LEU2 locus was confirmed by colony PCR using primers designed in the Yeast MoClo toolkit (*57*).

- LEU2-5’ forward CATAAATACCTTTCAAGC
- LEU2-5’ reverse TACAATCCTTGCCCGTGATG
- LEU2-3’ forward AGAGCACTTGAATCCACTGC
- LEU2-3’ reverse GATTTGGTTAGATTAGATATGGTTTC

Integration events were verified using a colony PCR method based on a formamide extraction approach (FEA) that I developed. Briefly, a small portion of a single yeast colony was resuspended in 70 µL FEA buffer (98% formamide, 10 mM EDTA), incubated at 70 °C for 10 min, vortexed for 1 min, diluted with 200 µL water, and centrifuged at >10,000 g for 1 min to pellet debris. The resulting supernatant, containing nucleic acids, was used directly as a PCR template. PCR was carried out with Q5 high-fidelity DNA polymerase, using ≤2% (v/v) diluted FEA/DNA mix per reaction to avoid polymerase inhibition. Thermocycling conditions included an initial denaturation at 98 °C for 30 s, followed by 40 cycles of 98 °C for 12 s, annealing at the appropriate primer-specific temperature for 10 s, and extension at 72 °C for a duration proportional to amplicon length, with a final extension at 72 °C for 2 min. Amplified products were resolved by agarose gel electrophoresis. Positive colonies showed the expected ∼500 bp band, while negative controls produced no detectable product.

### Fitness screening of Dsup-expressing cells

To assess the fitness impact of Dsup expression under different plasmid backbones, promoters, and induction conditions (dextrose, galactose, or estradiol at varying concentrations), cells were grown in 96-well plates and monitored using an Agilent BioTek LogPhase 600 plate reader. Cultures were incubated at 30 °C with shaking at 800 rpm, and optical density (OD) measurements were recorded every 20 min over 48–72 h.

For quantitative comparisons, cultures were inoculated at an initial OD600 of 0.06. Doubling time was calculated as the time elapsed from inoculation until the culture reached a five-generation point (OD600 ≈ 1), divided by five. Inhibition rates and visualization of growth curves were performed using AUDIT, an R package and online tool developed in our laboratory (*58*). For strains carrying centromeric or integrating plasmids with estradiol-inducible promoters, control cultures consisted of BY4741 transformed with the corresponding empty vector (identical plasmid lacking the Dsup-GFP or GFP ORF). Inhibition rates were plotted relative to this control to account for background plasmid effects.

### Protein expression dynamics via fluorescence microscopy and flow cytometry

To monitor expression of Dsup-GFP and GFP, we used fluorescence microscopy and flow cytometry. In the early stages of the study, fluorescence microscopy was used to confirm whether cell populations expressed the protein of interest. Samples were visualized on an EVOS fl AMF4300 microscope via the GFP channel (excitation 470/ emission 525nm).

To visualize GFP signals directly from transformation plates, we used a Sapphire FL Biomolecular Imager (only for centromeric plasmids of Dsup-GFP or GFP under the TDH3 promoter).

Quantification of relative expression levels was performed using a Beckman Coulter CytoFLEX LX flow cytometer. For each assay, BY4741 cells with the integrated cassette were inoculated into SC medium at starting OD of 0.06, distributed into 96-well plates (98 µL/well), and induced with 2 µL of estradiol diluted in 10% ethanol to achieve final concentrations ranging from 0 to 300 nM (and 0.02% ethanol). Each condition was assayed in triplicate. Cultures were grown at 30 °C with shaking at 800 rpm, and samples were collected for flow cytometry when all cultures reached mid-log phase (OD > 0.5). Cells were sonicated for 30 sec and analyzed immediately using the FITC channel without prior fixation. For each sample, ≥ 10,000 gated events were recorded, with gating set to exclude debris and small particles. Fluorescence values (geometric mean, FITC channel) were normalized between 0 and 100 prior to analysis. GraphPad Prism software (version 10.6.0) was used for data visualization and curve fitting. To determine the estradiol concentration at which expression reached a plateau, fluorescence intensities from three biological replicates were averaged and fit with a sigmoidal dose–response curve, as fluorescence typically follows a saturation profile with respect to ligand concentration (*59*). The gradient of the fitted curve was then examined, and the plateau region was defined as the concentration range where the rate of increase dropped below 0.5, indicating negligible further gain in fluorescence.

For quantification of protein expression from microscopy images, the total number of cells and the number of GFP-positive cells in each image were counted manually using an iPad and Apple Pencil.

### Cellular localization

For nuclear localization, yeast cells expressing Dsup-GFP at 75 nM estradiol were collected from log-phase cultures by centrifugation at 3000×g. The medium was discarded, and the cells were resuspended in PBS containing 0.01% Triton X-100 and incubated for 2 minutes before centrifugation. The pellet was then resuspended in 2 mL PBS supplemented with 6 µg/mL Hoechst 33258 (Sigma, 94403) and incubated in the dark for 30 minutes. After incubation, cells were pelleted at 3000×g, washed once with 1 mL PBS, and resuspended in 500 µL PBS. Samples were immediately subjected to microscopy using a Zeiss LSM 900 confocal microscope with 63x magnification.

For mitochondrial localization, cells carrying Cox4 tagged with mCherry and expressing Dsup-GFP under 75 nM estradiol induction were collected at log phase and washed once with PBS. Microscopy was performed immediately without staining, using a widefield microscope, Leica THUNDER 3D Cell Imager with 100x magnification.

### Chromatin silencing

We used two-compartment Petri dishes (Thomas Scientific, 1195W18) without estradiol on one side and with estradiol on the other side. The growth was tested in different conditions: SC, –Ura, SC+5FOA, –Ade, –Trp, each with and without 75 nM of estradiol. Cells were grown in two SC liquid cultures (0 nM and 75 nM) overnight. In the morning, while both cultures were in mid-log phase, the ODs were adjusted to 0.5. Both cultures were serially diluted in water with a 5x dilution factor and mixed properly. Then, 5 µL of Dsup^−^ cells was added to the non-inducing side of the plate, and the same volume of Dsup^+^ cells was plated to the inducing side of the plate. Plates were incubated at 30°C for ∼40 hours before images were captured.

### Oxidative stress survival test

2 cultures of Dsup-GFP were inoculated without and with 75nM of estradiol. When both cultures were in log phase, their ODs were adjusted to 0.5 in fresh SC medium, and 5 mM of hydrogen peroxide was added to both. Then cultures were incubated at 30°C with shaking at 220 rpm. After 30 min, both cultures were diluted 50 times in water. Next, 10 µL of each culture was diluted in 1 mL autoclaved water, and 500 µL of that was plated on YPD agar plates without estradiol. Plates were incubated at 30°C and photographed after ∼48 hours. The number of colonies was counted manually using an iPad. We then used GraphPad Prism (version 10.6.0) to t-test function to see if the number of colonies for Dsup^−^ and Dsup^+^ cells were statistically different. The two conditions each had 3 biological replicates, started from 3 single colonies.

### Apoptosis test

Yeast cultures were grown in SC medium containing either 0 nM or 75 nM β-estradiol. In the mid-log phase, the OD₆₀₀ of each culture was adjusted to 0.8. Hydrogen peroxide was then added to 1400µL of each culture to a final concentration of 3 mM, followed by incubation at 30 °C for 2 hours. Post-treatment, cells were pelleted at 7000×g for 3 minutes and washed once with 1 M sorbitol. For spheroplast preparation, cell walls were enzymatically digested by resuspending the pellet in 500 µL spheroplast buffer (1 M sorbitol, 50 mM Tris-HCl, pH 7.5, 5 mM 2-mercaptoethanol) containing 2 mg/mL Zymolyase (AMSBIO, Cat# 120491-1) without shaking, followed by incubation at 30 °C for 15 minutes.

Spheroplasts were then collected by centrifugation at 3000 x g for 3 minutes and gently washed once with 1 M sorbitol. The resulting spheroplasts were resuspended in 500 µL of staining solution containing 1x Annexin V binding buffer, 5 µL Annexin V-PE/Cyanine5 (Elabscience, Cat# E-CK-A125), and SYTOX Green (Thermo Fisher S7020) at a final concentration of 1 µM, and incubated at room temperature for 20 minutes. Following staining, cells were centrifuged at 3000 x g for 3 minutes, resuspended in 1 mL Annexin V binding buffer, and maintained on ice. Flow cytometry analysis was performed immediately thereafter.

### Flow cytometry for Rnr3-GFP measurements

For this experiment, Dsup was tagged with mCherry to enable simultaneous tracking of Dsup and Rnr3-GFP expression by flow cytometry. Two cultures of Dsup^−^ (0 nM estradiol) and Dsup^+^ (with 75 nM estradiol) were grown in SC medium. During mid-log phase, the ODs were adjusted at 0.5 in fresh SC medium. MMS, hydrogen peroxide, and HU were added to both populations at 0.015%, 0.0005%, and 15 mM, respectively. The incubation time was 5 hours for MMS and HU, and 10 hours for H2O2. After incubation, we proceeded to flow cytometry immediately. The background was adjusted using the WT BY4741 strain, and any FITC signal above that was considered positive GFP signal. Then the number of GFP^+^ cells in each culture (Dsup^−^ or Dsup^+^) was recorded, and GraphPad Prism (version 10.6.0) to t-test function t-test function was used to see whether or not they were statistically different. Flow cytometry was carried out using a BECKMAN CytoFLEX LX machine. Each treatment had four biological replicates.

### Cell cycle study via flow cytometry

Cell cycle analysis was carried out based on the protocol by Rosebrock (*60*). The BY4741 strain transformed with Dsup-Cherry was induced with 0 nM, 75 nM, or 200nM estradiol, grown in SC medium, and cells were fixed at OD ∼0.8 by adding two volumes of 95% ethanol directly to each culture, gently mixed, and incubated at −20°C overnight to ensure permeabilization. Fixed cells were then pelleted by centrifugation at 5000 × g for 20 min at 4°C, and the ethanol supernatant was removed. The pellets were resuspended in 800 μL of 50 mM sodium citrate buffer (pH 7.2), vortexed to mix, and incubated at room temperature for 10 min to ensure complete rehydration and ethanol removal. Cells were collected again by centrifugation at 5000×g for 5 min at room temperature, and the supernatant was discarded. This wash step was repeated once more. The resulting pellet was resuspended in 500 μL of 50 mM sodium citrate buffer (pH 7.2) containing 20 μg/mL RNase A and 2.5 μM SYTOX Green (Thermo Fisher S7020), vortexed, and incubated at 37°C for 90 min in the dark. After RNase digestion, 10 μL of 20 mg/mL proteinase K was added, and the sample was vortexed briefly and incubated for 2 h at 55°C in the dark. This step is essential for minimizing multicell aggregates and obtaining uniform optical scatter properties. Following digestion, cells were stored at 4°C overnight in the dark. The next day, prior to flow cytometric analysis, samples were transferred to appropriate tubes and sonicated with 5 pulses using a microtip probe sonicator to dissociate mother-daughter pairs and cellular aggregates. A total of 60,000 gated single-cell events were recorded per sample using a flow cytometer, and the data were analyzed using FlowJo 10.10.0. Each condition (0 nM or 75 nM, or 200nM) had two biological replicates.

### Sample collection for RNA-seq and MNase-seq

To maximize the relevance of comparison between gene expression and chromatin accessibility, samples for RNA-seq and MNase-seq were collected from the same cultures. Briefly, 40 mL of each culture was started at OD of 0.05 and grown in SC at 30°C overnight, and cells were collected at log-phase (OD ∼0.8). 50 OD units and 10 OD units of cells were collected for MNase-seq and RNA-seq, respectively. Samples without induction were grown with 0 nM estradiol, while induced samples were grown with 75 nM estradiol.

### RNA Extraction and Library Preparation for Sequencing

Total RNA was extracted from yeast following the protocol adapted from Shedlovskiy et al. (*61*) with modifications. For each extraction, 6-10 OD units of a yeast culture in SC were collected during log phase. Cells were centrifuged to remove the medium, resuspended, washed with 1 mL deionized water, transferred to an Eppendorf tube, and recentrifuged to discard all residual water. Next, 250 µL of FEA mix (98% formamide + 10 mM EDTA) was added to the cell pellet, and samples were heated at 70°C for 10 minutes without shaking. After vigorous vortexing for 1 min and centrifuging at 15,000×g, 200µL of the RNA-containing supernatant was carefully transferred to a fresh Eppendorf tube. To this, 700 µL of RNAzol (Sigma R4533) was added, followed by the addition of 400 µL nuclease-free water to precipitate DNA, proteins, and polysaccharides. This mixture was vortexed and allowed to stand at 4°C for 15 minutes before centrifugation at >12,000×g for 15 minutes at 4°C. The upper supernatant layer (light blue), containing RNA, was transferred to a fresh tube. RNA was precipitated by adding an equal volume of 100% isopropanol, vortexing and incubating at -20°C for 30 minutes, followed by centrifugation at 12,000×g for 10 minutes at 4°C. The resulting RNA pellet was washed twice with 500 µL of 75% cold ethanol, centrifuged at 12,000×g for 5 minutes, and the ethanol was removed thoroughly. Finally, the RNA pellet was air-dried at 40°C for 3–4 minutes and resuspended in 200 µL of nuclease-free water. Lastly, RNA concentration and quality were assessed using a Nanodrop spectrophotometer and Bioanalyzer. RNA sequencing libraries were prepared from 500 ng total RNA per sample using the TruSeq® Stranded mRNA Library Prep Kit (Illumina, Cat. #20020594) according to the manufacturer’s standard protocol. Briefly, polyA-containing mRNA was isolated, fragmented, and reverse transcribed to cDNA, followed by second strand synthesis, A-tailing, adapter ligation, and PCR amplification using indexed adapters as described in the manufacturer’s guidelines. Library quality and concentration were assessed prior to sequencing. Libraries were normalized, pooled, and sequenced using Illumina NovaSeq 6000 (Paired Read), targeting >10 million reads per sample.

### Sample and library preparation for MNase-seq

MNase digestion protocol was adapted from a protocol by McKnight *et al.* (*62*) with minor modifications. Cells (50 OD units) were collected at OD ∼0.8, pelleted at 10,000×g for 5 min, washed with ice-cold deionized water, and pelleted again. Pellets were immediately crosslinked with 1% formaldehyde for 20 min at 25°C with shaking at 500 rpm, and crosslinking was quenched with glycine (final concentration 125 mM), followed by another wash with ice-cold 1 M sorbitol and centrifugation at 10,000×g for 3 min. For spheroplasting, cells were carefully resuspended in 1.4 spheroplast buffer (1 M sorbitol, 50 mM Tris pH 7.5, 5 mM 2-mercaptoethanol, 2 mg/mL Zymolyase (AMSBIO 120491-1)) and incubated at 30°C for 45 min without shaking. After centrifuge at 4,000 × g for 5 min, spheroplasts were gently washed once with ice-cold 1 M sorbitol and resuspended in 150 µL of MNase digestion buffer (1 M sorbitol, 50 mM NaCl, 10 mM Tris pH 7.5, 5 mM MgCl₂, 1 mM CaCl₂, 0.075% NP-40, 0.5 mM spermidine, 1 mM 2-mercaptoethanol) containing 15 U Exonuclease III (NEB M0206) and 20 U Micrococcal Nuclease (ThermoFisher 88216), and digested at 37°C for 20 min. Digestion was stopped by adding 20 µL STOP buffer (EDTA 0.5 M and EGTA 0.5 M, pH 8), followed by the addition of 4 µL RNase A (20 mg/mL) and incubation at 42°C for 60 min. Subsequent protein digestion and crosslink reversal were performed by adding 17 µL 10% SDS and 20 µL Proteinase K (20 mg/mL) and incubating at 65°C for 90 min. DNA was extracted by adding 650 µL of DNAzol, followed by 400 µL of pure ethanol to precipitate the DNA. Samples were vortexed for 10-15 seconds, incubated for 10 min at –20°C, and centrifuged at 18,000×g to pellet the DNA. Pellets were washed twice by adding 0.9 mL of 75% ethanol, centrifuging between washes to remove impurities. After the final wash, DNA pellets were air-dried on a heat block set at 40°C temperature and resuspended in a minimal volume of 1× CutSmart buffer (∼40µL). To prevent self-religation of DNA fragments, 1 µL Quick CIP (NEB M0525) was added, and samples were incubated for 10 min at 37°C. Normalized amount of DNA samples were run on a 1.5% agarose gel at 150 V for 70 min; the mononucleosome band was excised and purified using a Qiagen MinElute column, with final elution in 20µL of 1× TE. NEBNext UltraExpress® DNA Library Prep Kit (E3325) and NEBNext® Multiplex Oligos for Illumina® (E6440) were used for sequencing library preparation following the manufacturer’s protocol. Libraries were normalized, pooled, and sequenced using Illumina NovaSeq 6000 (Paired Read), targeting 7-10 million reads per sample.

### SGA for generating the library of haploid deletions crossed with Dsup

The Query strain was constructed by transforming the SGA-starter strain (Y7092) with a LEU2-marked, integrating plasmid carrying an estradiol-inducible open reading frame. For SGA, the query strain was grown at 30 °C and the fresh culture was plated (1 mL of 0.2 OD per plate (Nunc™ OmniTray™ Single-Well Plate Thermo Fisher, 140156)) as lawns on SD–Leu agar, and incubated for 2 days at 30 °C. Mutant arrays were prepared from frozen 96-well 7% DMSO stocks of the yeast deletion (MATa xxxΔ::kanMX4). Plates were thawed at room temperature and pinned onto YPD+G418 agar plates using a Biomatrix Robot (S&P Robotics). Plates were incubated at 30 °C for 2 days. Four 96-format plates were condensed into a single 384-format plate using 96 pins and incubated for 2 days at 30 °C. Subsequently, query strains and mutant arrays were co-pinned onto YPD agar plates and incubated at 30 °C for 1 day to allow mating. Diploids were selected through two sequential rounds on SD–Leu+G418 plates (2 days each at 30 °C). Sporulation was carried out by transferring diploids onto sporulation medium supplemented with G418 and incubating at room temperature (°25 °C) for 7 days. Haploid MATα meiotic progeny engineered with pEstradiol-DSUP-GFP and a LEU2 marker were selected using two sequential rounds on SD–HRLK+Canavanine/Thialysine/G418 plates (2 days each at 30 °C). After the second round of selection, colonies were resuspended in sterile PBS buffer, pooled at OD600 of 40, and immediately frozen in 7% DMSO at -80 °C.

### Pooling the SGA final collection

Final SGA plates (–HRLK+can/thia/G418) were grown for two days at 30 °C. 8 mL of PBS was added to each plate, and colonies were resuspended using a sterile spreading stick. The suspension was quickly collected into an Erlenmeyer flask, and the OD600 was adjusted to 40. DMSO was added to the pooled culture to a final concentration of 7%. The pool was then aliquoted into smaller volumes and stored at –80 °C.

### Bar-seq library preparation

Bar-seq screens were carried out based on the standard protocol described by Barazandeh *et al.* (*63*). Pooled homozygous deletion strains previously crossed with the Dsup-expressing query strain via SGA were cultured in SC–HRUK, supplemented with canavanine, thialysine, and G418 to maintain selection pressure. Based on prior observations showing that Dsup expression at 75 nM estradiol and that growth in – HRUK medium amplified fitness defects, we optimized the induction condition to 40 nM estradiol. This concentration allowed for robust Dsup expression while maintaining an approximate 20% growth di3erence between Dsup– and Dsup+ cultures. Experimental cultures were inoculated at an initial OD600 of 0.0625 in 48-well plates and incubated at 30°C with continuous shaking, and DNA-damaging agents were applied at doses that produced 20–30% growth inhibition in the absence of Dsup induction. Growth was monitored by measuring OD600 every 20 minutes. An automated liquid handler (S&P Robotics, model GP2) was used to collect transfer cultures to 4°C at OD600 of 1 — equal to after 5 generations of growth. Genomic DNA was extracted from yeast cultures using the YeaStar Genomic DNA Kit (Zymo Research, D2002) according to the manufacturer’s instructions (chloroform-free approach). DNA concentration and purity were assessed with an Agilent Cytation 5 Take3 microplate reader. Bar-seq libraries were generated through a two-step PCR protocol. In the first round (PCR1), uptag and downtag barcode regions from the S. cerevisiae YKO deletion pool engineered with DSUP were independently amplified using common primers that flank Up tag and Down tag. PCR was carried out using a Phusion™ Hot Start II kit (Thermo Fisher, F549S) according to the manufacturer’s instructions. Thermocycling consisted of an initial denaturation at 98°C for 3 minutes, followed by 25 cycles of 98°C for 10 seconds, 59°C for 30 seconds, and 72°C for 20 seconds, with a final extension at 72°C for 5 minutes. Amplicon sizes were confirmed on a 1% agarose gel. Equimolar amounts of uptag and downtag products were pooled, diluted 10-fold, and used as templates for PCR2 to append dual-indexed Illumina Nextera adapters (i7/i5).

PCR2 was performed in a 50 µL reaction using a Phusion™ Hot Start II kit (Thermo Fisher, F549S) following standard protocol with 1.5 µL of diluted PCR1 product, and 0.5 µM of each PCR2 indexing primer. Cycling conditions were 98°C for 3 minutes, followed by 6 cycles of 98°C for 30 seconds, 60°C for 30 seconds, and 72°C for 30 seconds, with a final extension at 72°C for 5 minutes. Final libraries were purified using HighPrep™ PCR Clean-up beads (1.2X bead-to-sample ratio), quantified with the Qubit™ dsDNA HS Assay (Thermo Fisher, Q33231), and quality-checked using Agilent High Sensitivity D1000 ScreenTape. Libraries were normalized, pooled, and sequenced using Illumina NovaSeq 6000, targeting 5 million reads per sample.

Common Primers used in Bar-seq Library Preparation:

- Illumina PCR1-Uptag

`o` 5’ TCGTCGGCAGCGTCAGATGTGTATAAGAGACAGGATGTCCACGAGGTCTCT 3’
`o` 5’ GTCTCGTGGGCTCGGAGATGTGTATAAGAGACAGGTCGACCTGCAGCGTACG 3’
- Illumina PCR1-Downtag

`o` 5’ TCGTCGGCAGCGTCAGATGTGTATAAGAGACAGGAAAACGAGCTCGAATTCATCG 3’
`o` 5’ GTCTCGTGGGCTCGGAGATGTGTATAAGAGACAGCGGTGTCGGTCTCGTAG 3’

### RNA-seq quality control and mapping

Raw RNA-seq reads were processed using the nf-core/rnaseq pipeline (version 3.14.0). The pipeline was executed with all default parameters. The nf-core/rnaseq workflow provides a standardized and reproducible framework for RNA-seq analysis, including quality control, read trimming, alignment, quantification, and comprehensive reporting (*64*). Initial quality assessment of raw sequencing reads was performed with FastQC, followed by adapter and low-quality base trimming using Trim Galore. Reads were then aligned to the Saccharomyces cerevisiae reference genome with STAR, using the default two-pass alignment strategy. Transcript quantification was carried out with featureCounts, and expression was additionally estimated with Salmon in quasi-mapping mode. Post-alignment quality control included the generation of MultiQC summary reports, which integrated metrics across all stages of the workflow. The reference genome and corresponding gene annotation files were obtained from the Illumina iGenomes resource (Ensembl source, build R64-1-1). The genome FASTA and annotation GTF files provided by this collection were used consistently for alignment and quantification.

### DiPerential expression and functional enrichment analysis

Downstream analysis of the RNA-seq data was performed in R (version 4.5.0). Gene-level counts from Salmon, generated within the nf-core/rnaseq pipeline, were imported and processed using the tximport and DESeq2 packages (*65*, *66*). Genes with fewer than two total reads across all samples were excluded. Normalization and variance stabilization were applied using the rlog transformation to account for sequencing depth and improve comparability between samples. Differential expression analysis was conducted with DESeq2, using a design that compared induced Dsup expression against the uninduced control. Genes were considered significantly differentially expressed if they exhibited an absolute log2 fold change greater than 1 and an adjusted p-value (Benjamini–Hochberg correction) below 0.01. Results were visualized with a customized volcano plot, highlighting significantly upregulated and downregulated genes. To investigate functional consequences of gene expression changes, a Gene Ontology (GO) over-representation analysis was carried out separately for upregulated and downregulated genes. The clusterProfiler package was used with the org.Sc.sgd.db annotation database, focusing on the Biological Process (BP) ontology (*66*). For each direction of change, the top ten enriched GO terms were retained and visualized using dot plots. Enrichment significance was based on adjusted p-values (FDR < 0.05).

### MNase-seq quality control, mapping, and visualization

Adapters and low-quality bases were trimmed from paired end reads using Trimmomatic (v0.39), and post-trim quality was assessed with FastQC (v0.12.1) (*67*). Trimmed reads were aligned with BWA-MEM (v0.7.18) to *S. cerevisiae* reference genome. TSS coordinates were taken from a curated BED file. We retained primary alignments with MAPQ ≥ 30 and excluded unmapped, secondary, QC-fail, and supplementary records. PCR/optical duplicates were removed, producing de-duplicated, coordinate-sorted BAMs. BAMs were indexed and alignment statistics were summarized with SAMtools flagstat (v1.19.2) (*68*). Genome-wide nucleosome occupancy, positions, and fuzziness were computed with DANPOS2 (*69*) dpos (v2.2.2.) using a fixed read-density cutoff (--height 5) with P-value thresholding disabled (--pheight 0). Differential nucleosome occupancy runs were performed, and standard DANPOS outputs were used for +1 nucleosome and fuzziness metrics. For visualization, dyad-centered coverage was derived from de-duplicated BAMs with a 147-bp fragment shift, normalized to the genome-wide mean (excluding zero bins), Gaussian-smoothed (window = 3), and converted to bigWig. Per-condition replicate bigWigs were averaged with deepTools v3.5.6 (*70*) (bigwigCompare --operation mean, bin size = 5). Differential occupancy was calculated as log₂ ratios (default pseudocount = 1; zero bins skipped) for treatment vs WT and dose contrasts. Signal matrices were generated around TSSs in reference-point mode (bin size = 5). Genes were ordered by mean signal within the -200 bp NDR upstream of the TSS, and this order was propagated to full ±1 kb heatmaps to ensure comparability across conditions. Heatmaps and average profiles were plotted with deepTools (plotHeatmap, plotProfile), and differential matrices were clustered by k-means (k = 5).

### Metagene profiling

MNase-seq coverage profiles were generated to visualize the average MNase-seq signal distribution relative to transcription start sites (TSSs) across all annotated genes. For each condition, MNase-seq coverage matrices were prepared with rows corresponding to individual genes and columns corresponding to fixed-width bins (5 bp) spanning −1000 bp to +1000 bp relative to the TSS (400 bins total). Rows in which all bin values were zero were excluded from further analysis. For each bin position, the mean coverage across all retained genes was calculated. To estimate uncertainty in the mean signal, we performed bootstrap resampling of genes (500 iterations, with replacement) and calculated the mean profile for each bootstrap sample. The 2.5th and 97.5th percentiles of these bootstrap distributions at each position were taken as the lower and upper bounds of the 95% confidence interval (CI). The resulting mean profiles were smoothed using a Gaussian kernel (σ = 2 bins) to reduce noise. Profiles for the two conditions were plotted with shaded ribbons representing the 95% CI.

### Fuzziness analysis

Box plots were generated from the DANPOS-derived output, which reports per-locus nucleosome fuzziness_log₂FC and fuzziness_di3_FDR for Dsup– vs WT and Dsup+ vs WT; the input was restricted to loci within coding regions. For each comparison, loci were filtered by FDR < 0.05, the passing log₂FC values were reshaped to long format, and plots were produced in R/ggplot2 as box plots (boxes = interquartile range, bold line = median, whiskers = 1.5×IQR) with a dashed y = 0 reference line. Statistical significance for each distribution was assessed using a two-sided Wilcoxon signed-rank test vs 0 (μ = 0).

### Bar-seq data analysis

For Bar-seq datasets generated with paired-end sequencing, read pairs were first combined into single contiguous reads using BBMerge v. 8.82 from the BBTools/BBMap package. After this initial step, all libraries—regardless of whether they were sequenced as paired- or single-end—were processed with the same analytical workflow. In brief, single-end Bar-seq reads were trimmed to 50 bp using Trimmomatic v. 0.33 (*67*) and subsequently aligned against a yeast barcode reference database with BWA v. 0.7.12 (*71*). This reference was constructed from the barcode collection published by Pierce *et al.* (*72*), with barcode primer sequences appended to both sides of the uptag and downtag barcodes. Alignment files were filtered with SAMtools v. 1.2 (*68*), retaining only reads with mapping quality scores ≥30. Read counts for each library were then extracted with BEDTools v. 2.24 (*73*), and the results were consolidated into a count matrix using a custom Perl script. To ensure robustness, only tags with at least 50 counts across all control replicates were considered; uptag and downtag counts were then summed per strain, normalized, and subjected to statistical testing using the edgeR package v. 3.10.5 (*74*), following the approach previously described (*75*).

## Author Contributions

Conceptualization: H.K.G. and C.N.; Methodology: H.K.G. and C.N.; Investigation: H.K.G. and L.H.; Formal analysis: H.K.G., C.N., M.B., and G.G.; Validation: H.K.G.; Data curation: H.K.G.; Writing – original draft: H.K.G. and C.N.; Visualization: H.K.G. and M.B.; Supervision: C.N.; Funding acquisition: C.N. and G.G. L.H. contributed as undergraduate trainee and received training from C.N. and H.K.G.

## Acknowledgments

H.K.G. thanks Ed Grant and Vivien Measday for their generous support and guidance. H.K.G. also thanks John Rinn for making his online course on omics mapping and data analysis freely available through open access. We thank Matej Usaj and Adam Sanford for helpful guidance and Eleanor Campbell for her assistance in the experimental work. We are very grateful to Lorraine Pillus, Grant Brown, Chris Loewen, and Hilla Weinberg for generously sharing strains.

