## supplementary figures for "Friend and Foe: Genome-Wide Analysis of the Tardigrade Dsup Protein Expressed in Yeast Reveals Trade-offs of DNA Protection"

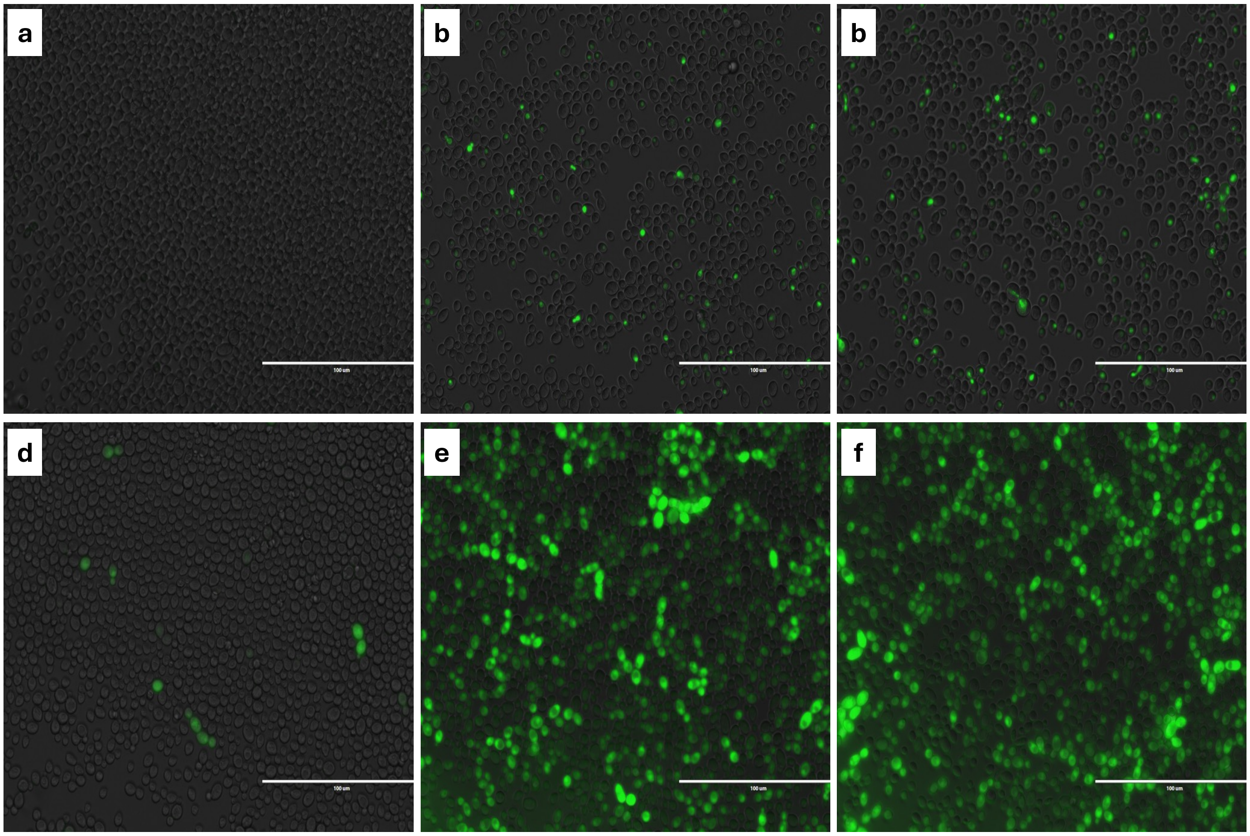


**Figure S1. Expression of Dsup and GFP using the CEN plasmid with the estradiol promoter.** (**a-c**) Dsup-GFP expression with 0, 50, and 100 nM of Estradiol. (**d-e**) GFP expression with 0, 50, and 100 nM of Estradiol.

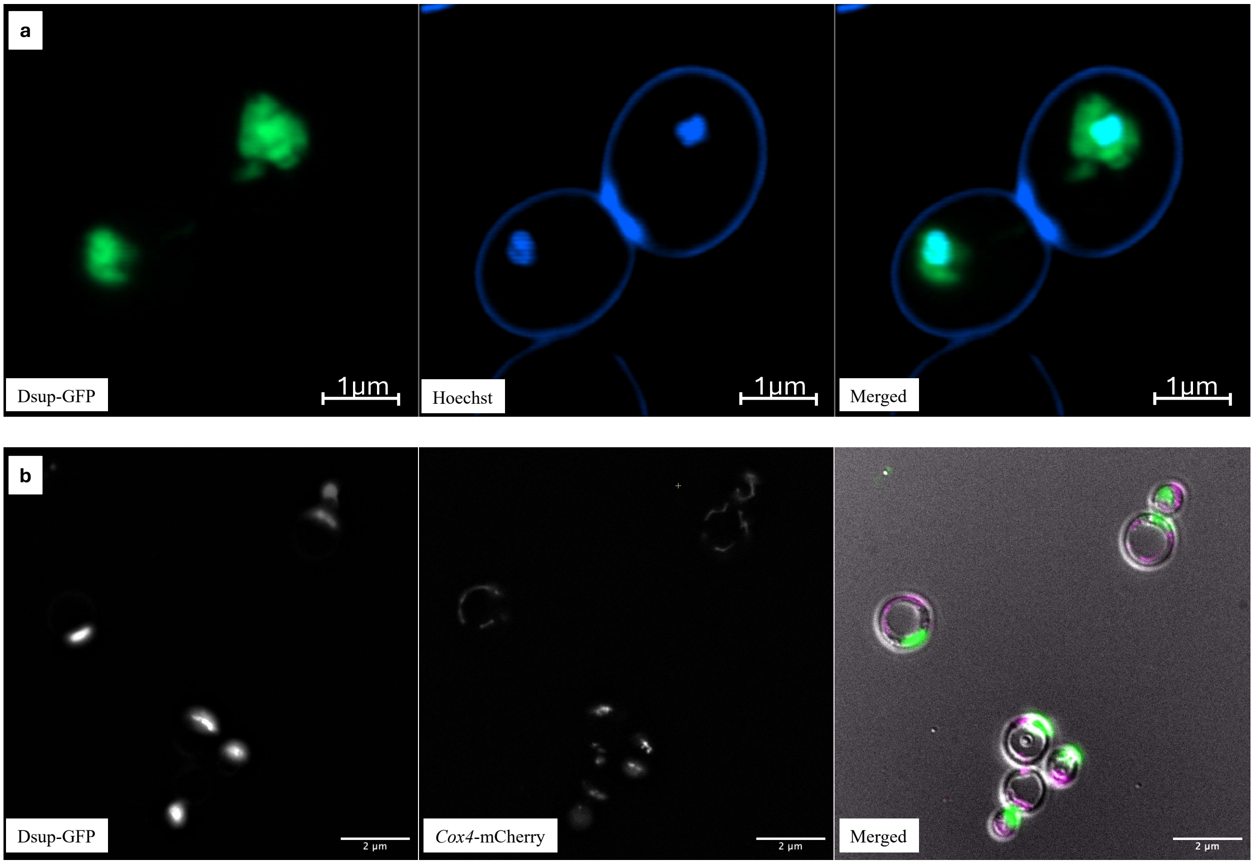


**Figure S2. Subcellular localization of Dsup in yeast.** (**a**) Dsup-GFP fluorescence (green) co-localizes with nuclear DNA stained by Hoechst (blue), confirming nuclear localization. Scale: 1μm (**b**) Co-expression of Dsup-GFP (green) and the mitochondrial marker Cox4-mCherry (magenta) shows no overlap, indicating that Dsup does not localize to mitochondria. Merged images include brightfield for visualization of cell morphology. Scale: 2μm

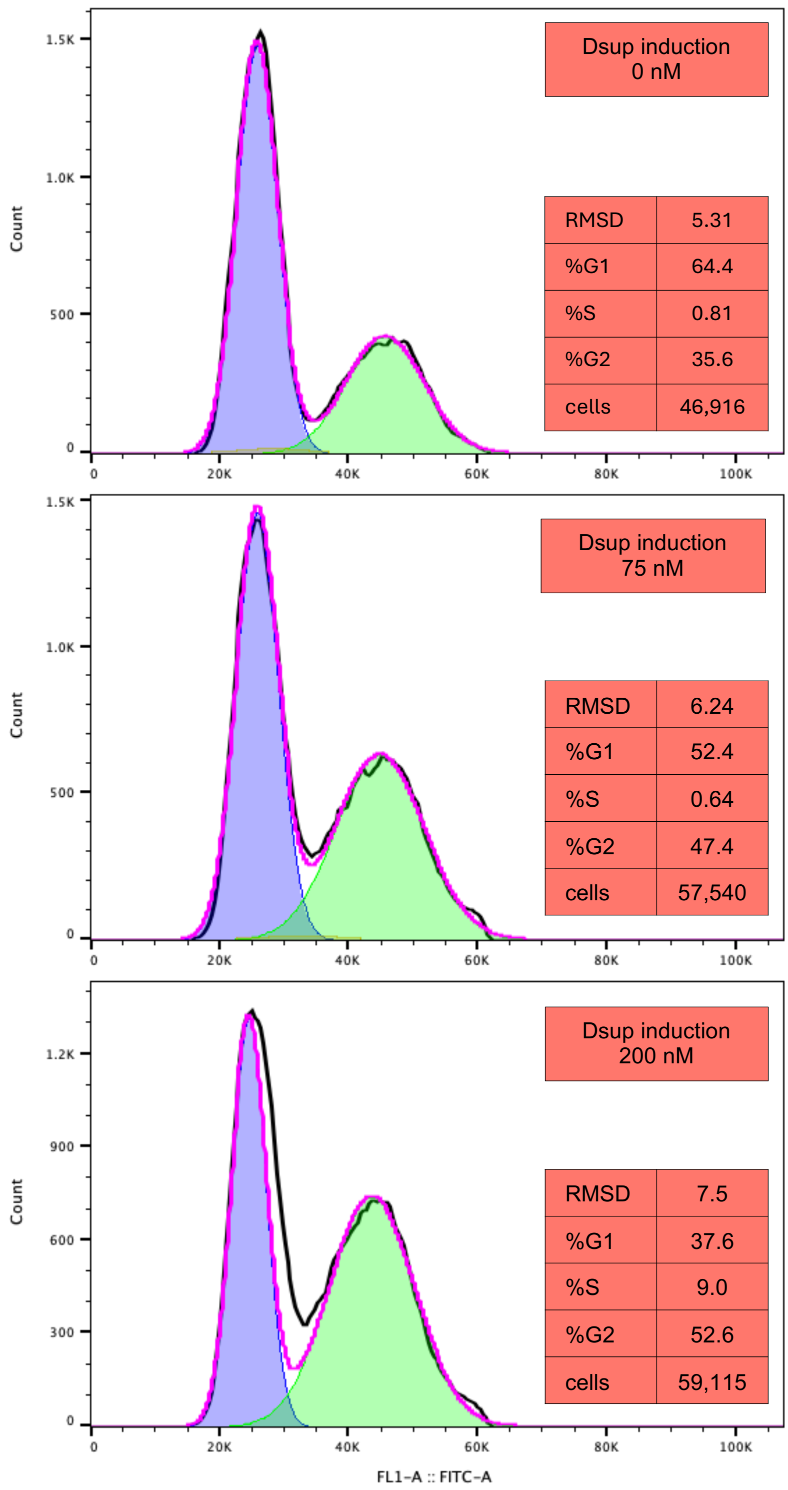


**Figure S3. Effect of Dsup expression on yeast cell cycle distribution.** Cell cycle profiles of yeast cultures transformed with the pEstradiol-Dsup at 0, 75, and 200 nM estradiol, with percentages of cells in G1, S, and G2/M shown in each plot. The black line shows the raw data, and the purple line is the cell cycle model automatically built using by FlowJo cell cycle function.
